# Selective class IIa HDAC inhibition reverses diastolic dysfunction in cardiometabolic HFpEF

**DOI:** 10.64898/2026.01.14.699428

**Authors:** Jiale Huang, Friederike Schreiter, Junyu Fan, Zihao Chen, Harikrishnareddy Paluvai, Coenraad Withaar, Christian U. Oeing, Ali Reza Saadatmand, Eric Schoger, Shaza Haydar, Adel Elsherbiny, Marco Hagenmüller, Karine Lapouge, Nuno Guimarães-Camboa, Inês Falcão-Pires, Nazha Hamdani, Almut Schulze, Sepp Jansen, Michael A. Nolan, Ralf Gilsbach, Rudolf A. de Boer, Matthias Dewenter, Johannes Backs

**Author notes:** Corresponding author: Johannes Backs, MD, Heidelberg University, Medical Faculty Heidelberg, Im Neuenheimer Feld 669, 69120 Heidelberg, Germany. These authors contributed equally (Jiale Huang, Friederike Schreiter, Matthias Dewenter, Johannes Backs).

## Abstract

Heart failure with preserved ejection fraction (HFpEF) is a highly prevalent cardiometabolic syndrome with yet no effective therapies. Here, we report that selectively class IIa histone deacetylases (HDACs) but no other classes of HDACs are enzymatically activated in hearts from HFpEF patients and cardiometabolic HFpEF animal models. Cell type-specific and enzymatic activity-specific genetic loss-of-function models of the cardiomyocyte-enriched class IIa HDAC family member **HDAC4** and the pharmacological class IIa HDAC-selective inhibitor TMP195 prevent and reverse diastolic dysfunction and exercise intolerance in cardiometabolic HFpEF *in vivo*. In contrast to pan-HDAC inhibition no adverse effects are observed. Despite its well-known non-enzymatic role as transcriptional repressor, we found that specifically enzymatic activation of HDAC4 has little direct effects on cardiomyocyte-intrinsic gene expression. Instead, non-epigenetic actions lead to endothelial activation via altered cardiocrine signaling. We discovered that selective enzymatic class IIa HDAC inhibition is a new therapeutic concept to combat cardiac HFpEF.

## INTRODUCTION

The prevalence of cardiometabolic diseases rises globally and treatment options are limited^1^. Heart failure with preserved ejection fraction (HFpEF) patients often present with a normal cardiac systolic function but an impaired diastolic function characterized by increased left ventricle (LV) filling pressure and prolonged LV relaxation^2,3^. Approximately half of HFpEF patients have major systemic comorbidities including obesity and hypertension and share a high risk for non-cardiovascular death compared to heart failure with reduced ejection fraction (HFrEF) patients^4^. Pathophysiological mechanisms of HFpEF differ substantially from HFrEF and many clinical trials have shown that there were no benefits for HFpEF patients by ‘canonical’ heart failure drugs such as betablockers and ACE inhibitors^5,6^. Notably, targeting metabolism systemically via sodium-glucose cotransporter 2 (SGLT2) inhibitors^7^ and Glucagon-like peptide-1 receptor (GLP-1) agonists^8^ demonstrated a significant reduction in risk of cardiovascular death, cardiovascular hospitalization and HF symptoms in HFpEF patients in recent studies. Both approaches, however, primarily affect non-cardiac cell types and therefore exert their favorable effects on the heart therefore in an indirect manner.

Recent studies on preclinical models of diastolic dysfunction have demonstrated that inhibition of histone deacetylases (HDACs) improves cardiac relaxation by affecting the acetylation of myofibrils and mitochondrial enzymes. These studies applied the HDAC inhibitors (HDACi) Givinostat (ITF2357)^9^ or SAHA (Vorinostat)^10,11^ which are non-selective or pan-HDAC inhibitors that block class I, class IIa and IIb HDACs^12^. Pan-HDAC inhibitors are approved for the treatment of multiple myeloma, T-cell lymphoma and, more recently, Duchenne muscular dystrophy, but their side-effect profiles including cardiovascular adverse effects does not qualify these pharmaceuticals for the long-term use in cardiovascular patients^13^. These adverse effects are attributed to inhibition of class I HDACs, which has been shown to cause arrhythmias, atherosclerosis and vascular calcification^12,14^. In contrast to class I HDACs, class IIa HDACs are best known as transcriptional repressors via their N-terminal domain, have much lower enzymatic activity on standard acetyl-lysine substrates and barely associate with histone tails^15–17^. We have previously shown that HDAC4, one class IIa HDAC, is cleaved in cardiomyocytes by the lipid droplet-associated protein ABHD5 to yield an N-terminal proteolytic fragment (HDAC4-NT), which protects the diabetic heart from failure^18,19^. Mechanistically, HDAC4-NT protects from systolic dysfunction by suppressing the hexosamine biosynthetic pathway and protein O-GlcNAcylation of Ca^2+^-handling proteins^20^. These studies revealed the critical non-enzymatic activity function of HDAC4 for cardiac contractility, while the role of the C-terminal enzymatic domain of HDAC4 remained elusive.

Lobera et al. developed a series of selective class IIa HDAC inhibitors based on a trifluoromethyloxadiazolyl (TFMO) moiety^21^. TMP195 (TFMO2) has been shown to enhance cancer therapy^22^, improve acute kidney injury^23^, blunt the increase of angiotensin II (Ang II)-induced arterial wall thickness^24^, and diminish plaque vulnerability^25^. While these data suggest a promising potential of class IIa HDAC inhibition in vascular pathology, protective effects on cardiac pathologies have not been shown.

In this study, we discovered that specifically the enzymatic activity of class IIa HDACs is increased in cardiometabolic HFpEF, and that enzymatic activation of HDAC4 in cardiomyocytes is causative for diastolic dysfunction via cardiomyocyte-endothelial cell crosstalk. Our results indicate that selectively targeting class IIa HDACs and in particular HDAC4 is a promising novel approach to treat HFpEF.

## RESULTS

### Class IIa HDAC activity is increased in HFpEF hearts

To explore the enzymatic activity of the different HDACs in HFpEF, we performed class-selective HDAC activity assays cardiac tissue and found that cardiac class IIa HDAC activity was increased in HFpEF hearts from mice subjected to high-fat diet (HFD) + L-NAME and in aged female mice subjected to HFD + AngII. However, neither class I nor class IIb HDAC activity showed any significant differences (Fig. 1a-f). We also observed a significant increase of cardiac class IIa HDAC activity but not class I or class IIb HDAC activity in HFpEF rat hearts (ZSF1 obese) compared the control rats (ZSF1 lean) (Fig. 1g-i). In myocardial biopsies from HFpEF patients, class IIa but not class I HDAC activity was elevated in many (but not all) patients compared to HFrEF patients (Fig. 1j, k). The obtained material from the respective biopsies was not sufficient to measure class IIb HDAC activity. These HFpEF patients were characterized by increased left ventricular peak systolic and end-diastolic pressure (LVPSP and LVEDP, respectively) while gender distribution, age or heart rate (HR) were not significantly different between HFpEF and HFrEF patients (Fig. 1l).

**Fig. 1.**
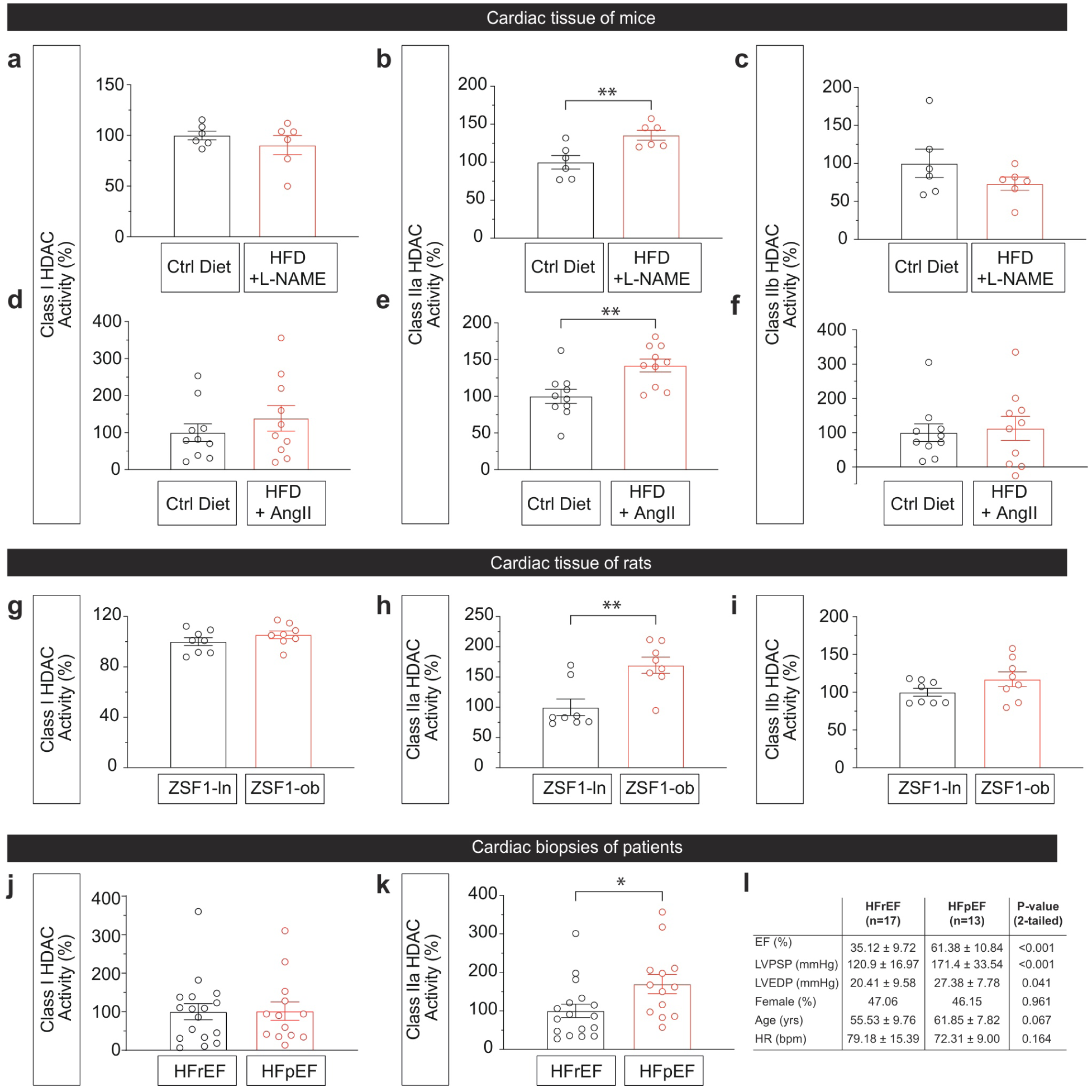
Cardiac HDAC activity in mice, rats and patients with HFpEF. **a-c,** Class I (a), class IIa (b) or class IIb (c) HDAC activity of cardiac tissue from HFD + L-NAME induced HFpEF mice or control mice. n=6/ group. **d-f,** Class I (d), class IIa (e) or class IIb (f) HDACs activity in HFD+ AngII induced HFpEF aged female mice or control mice. n=10/ group. **g-i,** Class I (g), class IIa (h) or class IIb (i) HDACs activity of cardiac tissue from ZSF1 obese (HFpEF rats) or ZSF1 lean rats (control rats). n=8/ group. **j, k,** Class I (j) and Class IIa (k) HDACs activity of cardiac biopsies from patients. n=17 in HFrEF group, n=13 in HFpEF group. **l,** Clinical characteristics of patients with HFpEF and HFrEF. Data are mean ± s.e.m. Statistical analysis was done by two-tailed unpaired t-test. ns = not significant; *P<0.05; **P<0.01.

### Cardiomyocyte HDAC4 contributes to diastolic dysfunction

Based on analysis of publicly available murine single-nucleus RNA sequencing (snRNA-seq) datasets^26^, *Hdac4* emerged as the most highly enriched class IIa HDAC in cardiomyocytes (CMs) (Extended Data Fig. 1a-d). On the basis of this observation, we investigated the potential causal role of HDAC4 in HFpEF using a mouse line with a cardiomyocyte-specific *Hdac4* knockout (cKO)^20^. In isolated adult CMs from cKO mice, total class IIa HDAC activity was reduced by approximately 50% compared with controls (Fig. 2a), confirming that *Hdac4* accounts for a major fraction of class IIa HDAC enzymatic activity in cardiomyocytes. The residual class IIa HDAC activity is likely attributable to other family members, including HDAC5, HDAC7 or HDAC9.

**Fig. 2.**
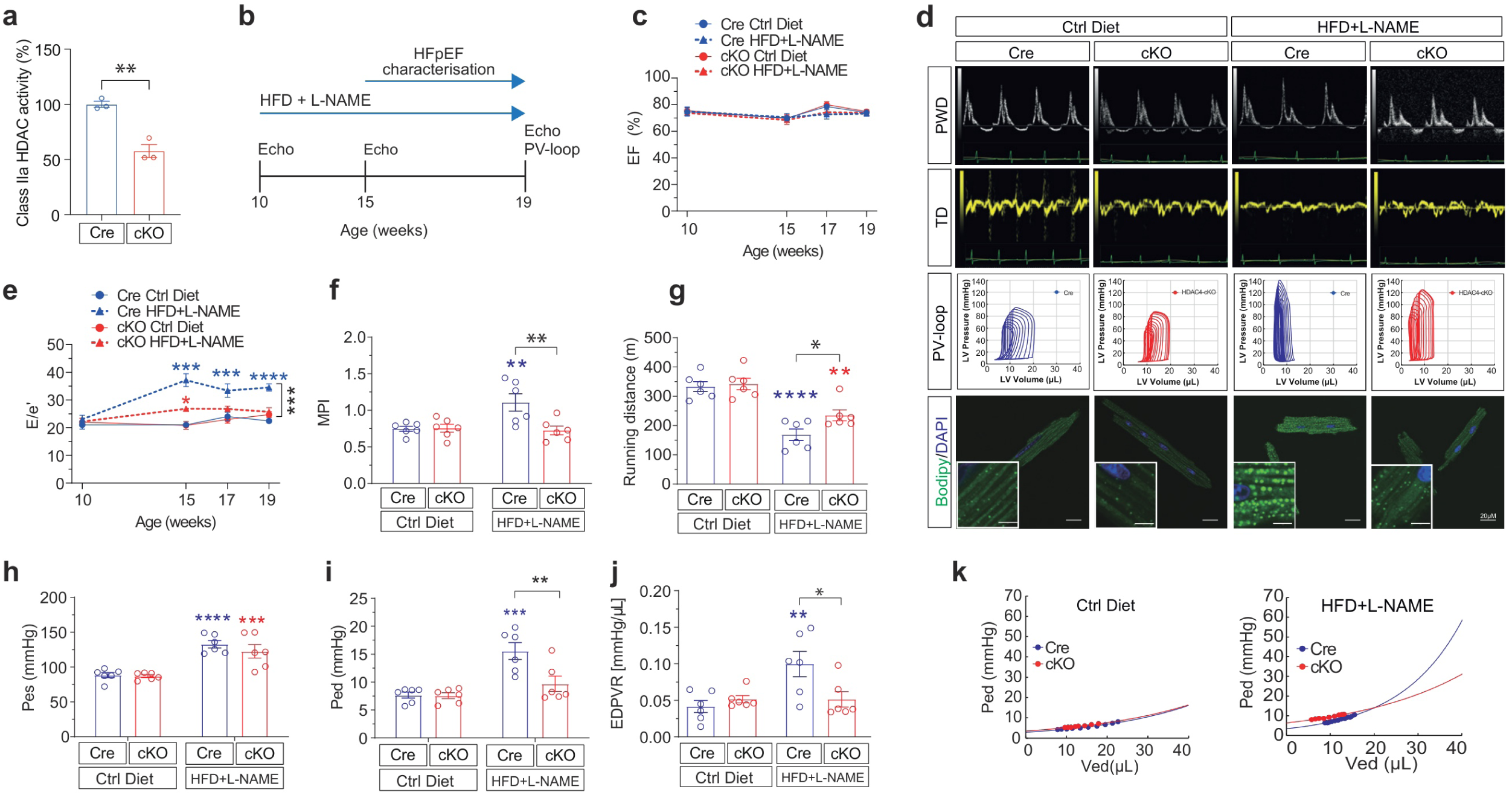
Cardiac HDAC4 is required for HFD + L-NAME induced HFpEF. **a,** Class IIa HDAC activity in living adult cardiomyocytes isolated from Cre or *Hdac4*-cKO (cKO) hearts. n=3/ group. **b,** Schema of the two-hit HFpEF mouse model. Mice were fed with HFD + L-NAME or Ctrl Diet during the whole period. HFpEF characteristics were evaluated after 5-week of different diets. **c,** Percentage EF of Cre or cKO mice with different diets. n=6/ group. **d,** Representative images from echocardiographic pulsed wave Doppler (PWD), TD (tissue Doppler), P-V loop (pressure-volume loop) and bodipy staining in different groups. **e,** The ratio between E wave and e’ wave of Cre or cKO mice with different diets. n=6/ group. **f,** Myocardial performance index (MPI) in week 19 of Cre or cKO with HFD + L-NAME or ctrl diet. n=6/ group. **g,** Running distance of mice during exercise exhaust test. n=6/ group. **h-j,** Left ventricle Pes (h), Ped (i) and slope of EDPVR (j) in Cre and cKO mice in response to HFD + L-NAME. Pes, end-systolic pressure; Ped, end-diastolic pressure; EDPVR, end-diastolic pressure-volume relationship. n=6/ group. **k,** Representative EDPVR exponential curve during hemodynamic measurements in mice with or without HFD + L-NAME. Ped, end-diastolic pressure; Ved, end-diastolic volume. Data are mean ± s.e.m. **a,** two-tailed unpaired t-test. **c, e,** two-way ANOVA with Tukey’s multiple comparisons test. **f-j,** two-way ANOVA with Sidak’s multiple-comparisons test. **\***P<0.05; **P<0.01; ***P<0.001; ****P<0.0001.

Cre-positive control mice (Cre) or cKO mice were subjected to HFD + L-NAME to induce HFpEF and its complications (Fig. 2b). While ejection fraction (EF) was preserved in all groups, we observed a significant increase of E/e’ after 5 weeks, 7 weeks and 9 weeks challenged with HFD + L-NAME only in Cre mice but not in cKO mice (Fig. 2c-e). Moreover, cKO mice showed no increases in myocardial performance index (MPI) (Fig. 2f), slope of end-diastolic pressure-volume relationship (EDPVR) and LV end-diastolic pressure (Ped) (Fig. 2h-k) which were observed in Cre mice after exposure to HFD + L-NAME. Microscopically, lipid droplet (LD) accumulation was found in adult cardiomyocytes from Cre but not *Hdac4*-cKO mice after HFD + L-NAME (Fig. 2d, Extended Data Fig. 2f). Exercise capacity was significantly decreased in Cre mice as an extra-cardiac manifestation of HFpEF, while a significant improvement of exercise capacity was significantly improved in cKO mice (Fig. 2g). Other systemic phenotypes including body weight, blood pressure, glucose tolerance test and the ratio of heart weight and tibial length showed no significance between Cre or cKO groups (Extended Data Fig. 2a-e). These results indicated that *Hdac4* in cardiomyocytes is essential for the development of diastolic dysfunction and exercise intolerance in HFpEF induced by HFD + L-NAME.

### The enzymatic activity of *Hdac4* contributes to diastolic dysfunction

To investigate the enzymatic activity of HDAC4, we generated an enzymatic class IIa loss-of-function HDAC4*^H976Y^* mutant (His-976-Tyr mutation, analogue to mouse His-968-Tyr). Baculoviral expressed recombinant HDAC4 mutant protein shows a significant reduction (>50-fold) of class IIa HDAC activity, but no difference in binding to class I HDACs (HDAC1, HDAC2 and HDAC3) (Fig. 3a, b) and the same suppression of myocyte enhancer factor 2 (MEF2) activity as HDAC4 wild type (WT) (Fig. 3c). Mice carrying *H968Y* mutation in HDAC4 (Fig. 3d, Extended Data Fig. 3a-c) show a 50% reduction in endogenous cardiac class IIa HDAC activity in the heart (Fig. 3e), confirming that most of the class IIa HDAC activity stems from HDAC4.

**Fig. 3.**
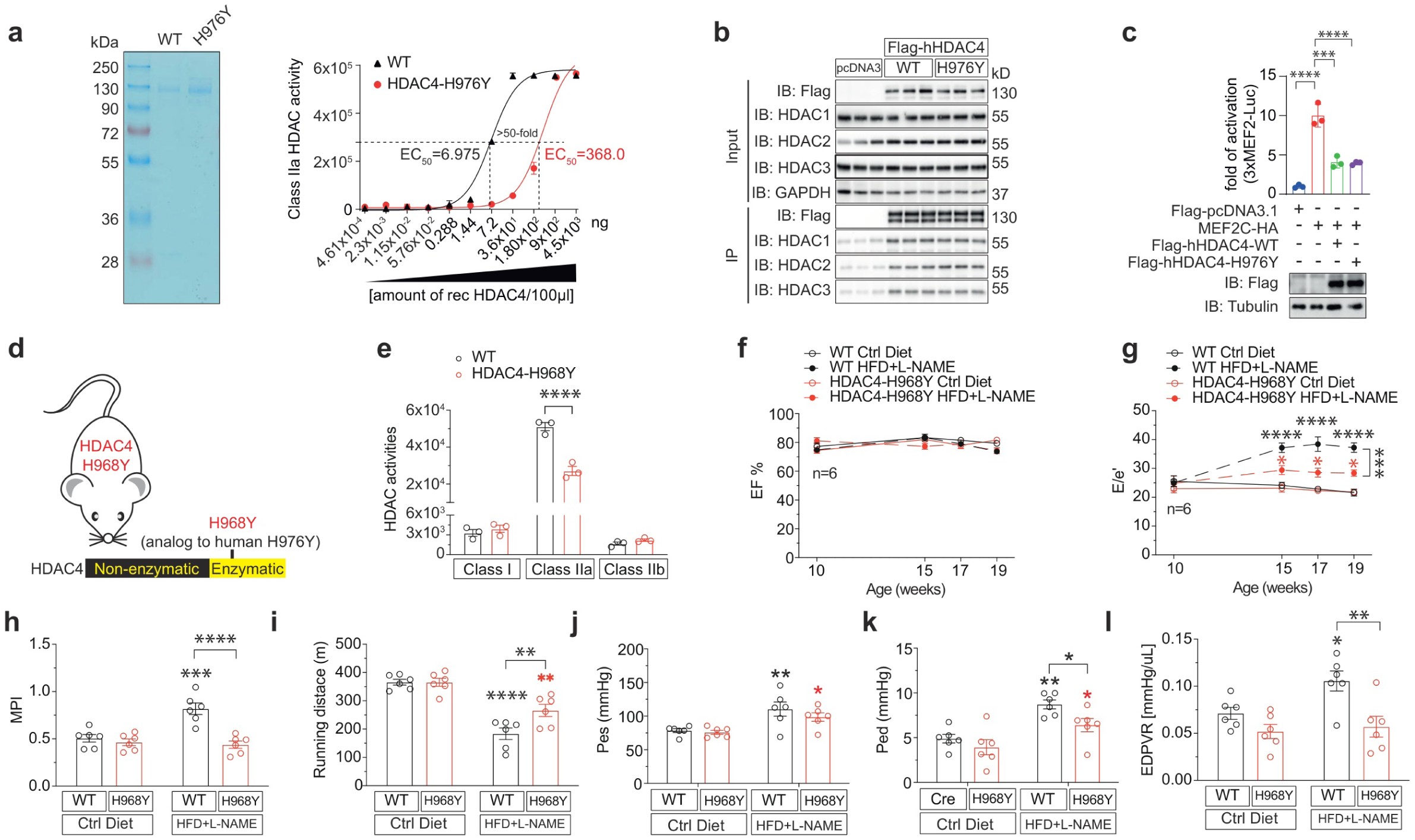
The enzymatic activity of HDAC4 is required for HFpEF. **a,** Coomassie-stained gel (left) and class IIa HDAC activity (right) of recombinant HDAC4 proteins (recHDAC4) from WT and H976Y mutant of human HDAC4 (H976Y). **b,** Immunoprecipitation on overexpressed Flag-HDAC4 WT and H976Y mutant in HEK293A cells. **c,** MEF2 luciferase activity on overexpressed HDAC4 WT and H976Y mutant in HEK293A cells. **d,** Schema of the strategy to generate *Hdac4^H968Y^* mouse line. **e,** Cardiac HDAC activity in *Hdac4^H968Y^* and WT mice. n=3/ group. **f, g,** Percentage of EF (f) and E/e’ (g) of week 10, 15, 17 and 19 of different diets in WT or *Hdac4^H968Y^* mice. n=6/ group. **h,** MPI of mice with or without 9-week HFD + L-NAME treatment. **i,** Running distance of exercise exhaustion test. n=6/ group. **j-l,** Pes (j), Ped (k) and EDPVR (l) in WT and *H968Y* mice in response to HFD + L-NAME. n=6/ group. Data are mean ± s.e.m. **c,** one-way ANOVA with Tukey’s multiple comparisons test. **e-g,** two-way ANOVA with Tukey’s multiple comparisons test. **h-l,** two-way ANOVA with Sidak’s multiple-comparisons test. **\***P<0.05; **P<0.01; ***P<0.001; ****P<0.0001.

To investigate the causative role of the enzymatic activity of HDAC4 in HFpEF, *Hdac4^H968Y^* mice were challenged with HFD + L-NAME. In contrast to WT littermates, *Hdac4^H968Y^* mice were largely protected from developing HFpEF, as confirmed by echocardiographic, hemodynamic measurements and preserved exercise tolerance (Fig. 3f-j, Extended Data Fig. 3i, j). Furthermore, *Hdac4^H968Y^*mice presented less LD accumulation in cardiomyocytes (Extended Data Fig. 3i, k). Notably, no significant difference in body weight, blood pressure, glucose tolerance and heart weight were observed between WT and *Hdac4^H968Y^* under HFD + L-NAME (Extended Data Fig. 3d-h). Taken together with the cKO-derived results, our data suggests that reduction of class IIa HDACs enzymatic activity in cardiomyocytes largely protects from the development of diastolic dysfunction and exercise intolerance in HFpEF.

To explore whether the enzymatic activity of HDAC4 is specifically involved in the development of HFpEF, we subjected the *Hdac4^H968Y^* mutant mice to transverse aortic constriction (TAC)-induced HFrEF. *Hdac4^H968Y^*mice developed the same degree of cardiac hypertrophy and contractile failure 8 weeks after TAC as WT littermates (Extended Data Fig. 3l-o), indicating that the enzymatic activity of HDAC4 is a critical driver of HFpEF but not HFrEF.

### TMP195 selectively inhibits class IIa HDACs

TMP195 selectively inhibited class IIa HDACs activity of recombinant HDAC4 protein, induced pluripotent stem cells (iPSCs)-derived cardiomyocytes or cardiac tissue, whereas class I or class IIb HDAC activity was not affected in HDAC activity assays (Extended Data Fig. 4a-c). In addition, we performed the assay on different amounts of murine cardiac lysates from WT and *Hdac4^H968Y^*with or without 3μM TMP195 to confirm the selective class IIa HDAC inhibition of TMP195 (Extended Data Fig. 4d-f). On top of this *ex vivo* data, we also tested TMP195 on a series of concentrations of recombinant human WT HDAC4 (recHDAC4) and the H976Y mutant (recHDAC4*^H976Y^*). 3μM TMP195 significantly inhibited class IIa HDAC activity of WT HDAC4, but not of recHDAC4*^H976Y^*(Extended Data Fig. 4g-i). Next, we tested the *in vivo* effects of the selective class IIa HDACi (TMP195 and TMP269) and pan-HDACi (SAHA). We found a significant decrease of class IIa HDACs activity in cardiac lysates after 4 hours of injection with TMP195 (15 mg/kg) or SAHA (10 mg/kg) (Extended Data Fig. 4j).

### Class IIa HDACs inhibition reverses HFpEF

To investigate the preventive effects of HDACi during the development of HFpEF, mice were pretreated with HFD + L-NAME one week prior to daily treatment (i.p./q.d.) with the selective class IIa HDAC inhibitor TMP195 or the pan-HDAC inhibitor SAHA for four weeks (Fig. 4a). TMP195 as well as SAHA reversed E/e’ to baseline levels after 2 weeks of treatment (Fig. 4b, c, Extended Data Fig. 5d). Also increases in MPI and EDPVR were completely reversed by TMP195 as well as SAHA (Fig. 4d, e, Extended Data Fig. 5d, e). Of note, TMP195 treatment significantly improved exercise capacity under HFD + L-NAME treatment up to the levels of control diet mice, whereas SAHA only showed a slight trend towards improvement of exercise tolerance (Fig. 4f). Neither TMP195 nor SAHA had significant effects on blood pressure and glucose tolerance (Extended Data Fig. 5a-c). We detected the expression of pro-fibrotic genes *Col1a1* and *Col3a1* induced by HFD + L-NAME while TMP195 completely prevented the upregulation of pro-fibrotic genes and SAHA had no significant effect (Extended Data Fig. 5f).

**Fig. 4.**
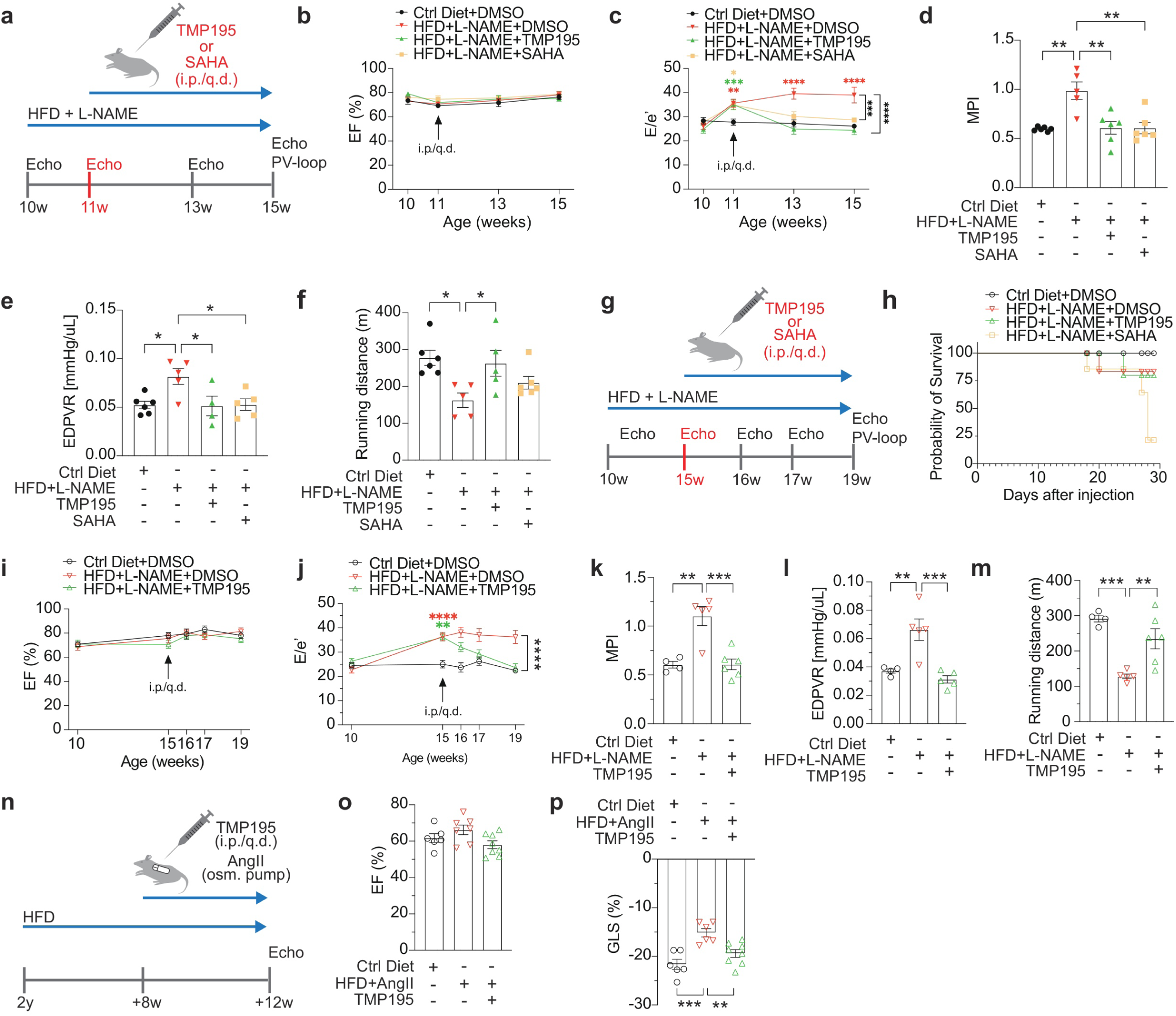
Early or late interventions with HDAC inhibitors reverse diastolic dysfunction in mice. **a,** Experimental design. Mice were prioritized treated with HFD + L-NAME for 1 week, DMSO (vehicle), TMP195 (15mg/kg) or SAHA (10mg/kg) was daily intraperitoneal injection (i.p./q.d.) for four weeks. Ctrl diet or HFD + L-NAME exposure was maintained for the whole period. **b, c,** EF percentage (b) and E/e’ (c) of week 10, 11, 13 and 15 with DMSO, TMP195 or SAHA treatment. n=6/ group except n=5 in HFD + L-NAME + DMSO group. **d-f,** MPI (d), slope of EDPVR (e) and running distance of exercise exhaustion test (f) in mice with different treatments. **g,** Experimental design. Mice were prioritized treated with HFD + L-NAME for 5 weeks, DMSO (vehicle), TMP195 (15mg/kg) or SAHA (10mg/kg) were daily intraperitoneal injection (i.p./q.d.) for four weeks. Ctrl diet or HFD + L-NAME maintained for 9 weeks. **h,** Probability of survival after daily injection. At “0” time point, n=4 in Ctrl Diet+ DMSO group, n=6 in HFD + L-NAME + DMSO and HFD + L-NAME + TMP195 group, n=5 in HFD + L-NAME + SAHA group. **i, j,** EF percentage and E/e’ of week 10 (5 weeks ahead of injection), 15, 16, 17 and 19 with DMSO or TMP195. n=4 in Ctrl Diet+ DMSO group; n=5 in HFD + L-NAME + DMSO; n=6 in HFD + L-NAME + TMP195 group. **k-m,** MPI (k), slope of EDPVR (l) and running distance of exercise exhaustion test (m) in mice with different treatments. **n,** Experimental design in aged mice. AngII-mini pumps were implanted in mice for 4 weeks after 8-week HFD treatment in 2-year-old (2y) female mice. TMP195 was daily intraperitoneal injection (i.p./ q.d.) after 8 weeks of HFD exposure. **o, p,** Percentage of EF (o) and percentage of GLS (p) in different groups. Data are mean ± s.e.m. **b, c, i, j,** two-way ANOVA with Sidak’s multiple-comparisons test. **d-f, k-m, o, p,** one-way ANOVA with Tukey’s multiple comparisons test. *P<0.05; **P<0.01; ***P<0.001; ****P<0.0001.

To investigate the therapeutic effects of HDACi in HFpEF, we set up a late intervention study with TMP195 and SAHA in mice after 5 weeks of exposure to HFD + L-NAME (Fig. 4g). Concerningly, SAHA caused a dramatic mortality within 4 weeks of treatment compared to any other group (Fig. 4h), indicating toxic effects of long-term pan-HDACi treatment in the setting of established cardiometabolic disease. Subsequently, the SAHA group was excluded in the phenotyping panel at the end of the study. We observed preserved EF in all groups while HFD + L-NAME-treated mice with TMP195 showed a reversal of diastolic dysfunction when treated with TMP195, as determined by measuring E/e’, MPI and EDPVR (Fig. 4i-l, Extended Data Fig. 6d, e). TMP195 again significantly improved exercise tolerance under HFpEF conditions (Fig. 4m). No significant effects on blood pressure and glucose tolerance observed (Extended Data Fig. 5a, b) while the ratio of heart weight to tibial length ratio decreased after TMP195 treatment compared to HFD + L-NAME-treated group (Extended Data Fig. 6c). Next, we tested the therapeutic effect of TMP195 in another HFpEF mouse model^27^ where HFpEF was induced by HFD + AngII in aged female mice (Fig. 4n). Indeed, we found that global longitudinal strain (GLS) was significantly increased after 4 weeks of TMP195 treatment while EF was preserved in all groups (Fig. 4o, p). In addition, *Col1a1*, *Col3a1*, *Nppa* and *Gdf-15* were significantly downregulated by TMP195 treatment (Extended Data Fig. 6f).

In summary, these findings demonstrate that class IIa HDAC inhibition rescues the cardiac HFpEF phenotypes *in vivo*, without the obvious safety liabilities of pan-HDACi.

### A cardiomyocyte-intrinsic HDAC4-dependent mechanism in cardiomyocyte-endothelial cell crosstalk

To understand the role of enzymatic activation of HDAC4 in cardiac gene regulation during HFpEF, we first performed bulk RNA-sequencing on cardiomyocyte nuclei isolated from left ventricular tissue from HFD + L-NAME treated mice as described previously^28–30^. Surprisingly, this analysis revealed only a few differentially regulated genes upon HFD + L-NAME in WT mice. Differential gene expression (DEG) analysis identified 156 genes. Among these, 36 were decreased and 41 increased in both genotypes after HFD + L-NAME treatment (Extended Data Fig. 7a-c). Even more surprisingly, despite the phenotypic rescue we did not find clear *Hdac4*-dependent transcriptional changes within CM nuclei (Fig. 5a, b). Because the functional rescue is also seen in cKO mice, when *Hdac4* is only deleted in cardiomyocytes, this provokes the notion that the mode of action of enzymatic HDAC4 activation is gene expression independent. Consistently, *Hdac4^H968Y^* cardiomyocytes do not show any changes in chromatin accessibility (Fig. 5c, Extended Data Fig. 7d). To reveal that the enzymatic mode of action of class I and class IIa HDACs is indeed distinct with respect to chromatin accessibility, we treated human iPSC-derived cardiomyocytes (iPSC-CM) with the pan-HDAC inhibitor (TSA), the class I HDAC selective inhibitor (Romidepsin) and the class IIa HDAC selective inhibitor (TMP195). Strikingly, class IIa HDAC inhibition did not increase chromatin accessibility in stark contrast to class I HDAC inhibition (Fig. 5d), pointing to a clearly distinct mode of action of class IIa versus class I HDACs.

**Fig. 5.**
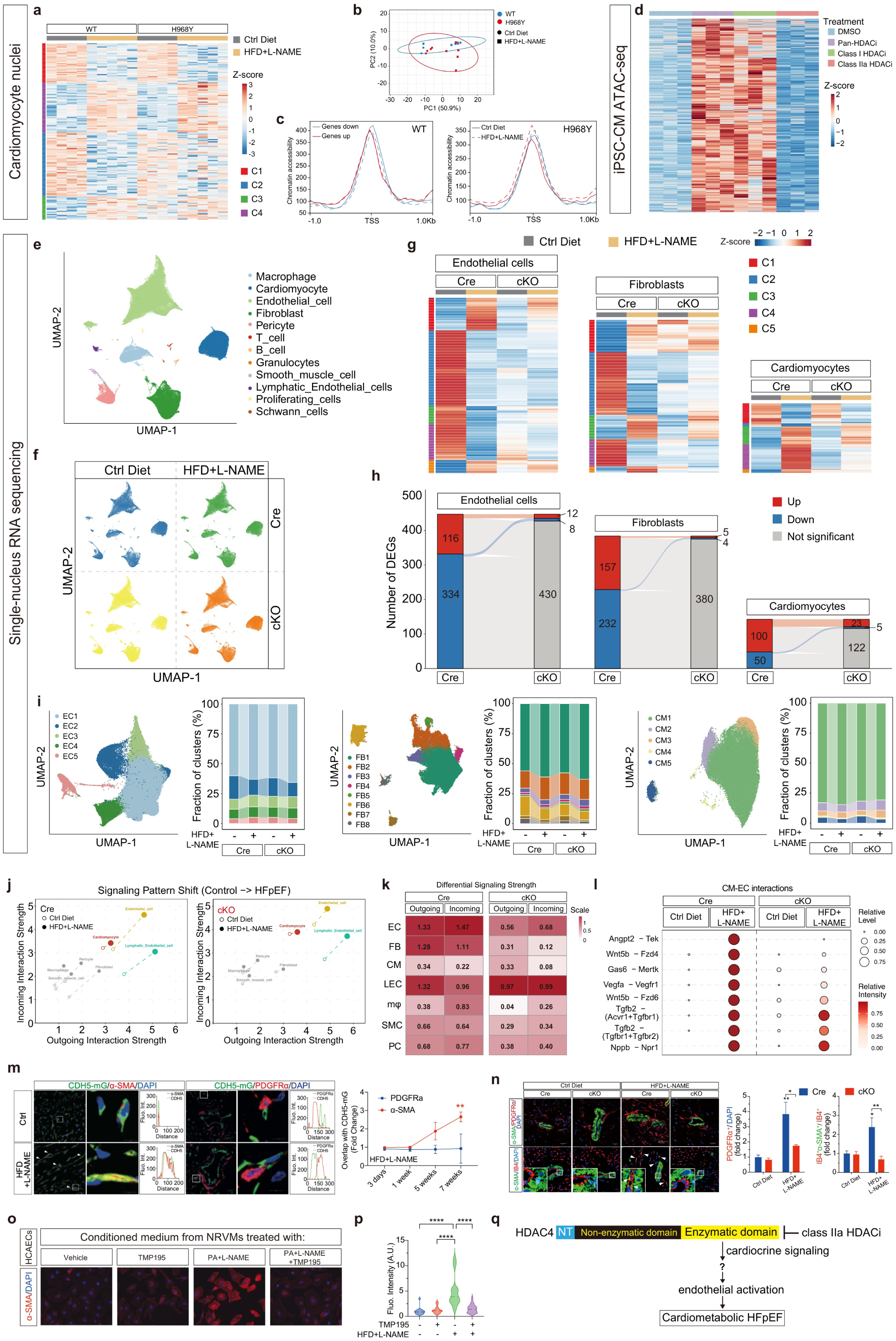
An HDAC4-dependent mechanism in cardiomyocyte-endothelial cell crosstalk. **a,** Heatmap of the differentially expressed genes (DEGs) responsive to the HFD + L-NAME diet, revealed by RNA sequencing of CM nuclei. n=4 in Ctrl Diet group, n=5 in HFD + L-NAME group. **b,** Principal component analysis (PCA) of WT or H968Y mice with or without HFD + L-NAME. n=4 in Ctrl Diet group, n=5 in HFD + L-NAME group. **c,** Average plots of chromatin accessibility (ATAC-seq) of peaks overlapping transcription start site of DEGs of HFD + L-NAME-responsive genes in WT and H967Y mutant. n=5 in Ctrl Diet group, n=6 in HFD + L-NAME group. **d,** ATAC-seq analysis for human iPSC-derived cardiomyocytes (iPSC-CM) treated with DMSO (Ctrl), the pan-HDAC inhibitor (TSA), the class I HDAC selective inhibitor (Romidepsin) and the class IIa HDAC selective inhibitor (TMP195). n=3 per group. **e, f,** Uniform manifold approximation and projection (UMAP) profiled by single-nucleus RNA sequencing uncovered 12 distinct cellular profiles (e) and distributions in different groups (f). n=3 per group. **g, h,** Heat map (g) and pseudobulk DEGs analysis (h) of endothelial cells (ECs), fibroblasts (FBs) and cardiomyocytes (CMs) from single-nucleus RNA sequencing. **i,** UMAP profiles and fractions of subclusters from ECs, FBs and CMs in different groups. n=3 per group. **j,** Trajectory analysis of the aggregate cell-cell communication network in Cre (left) and cKO (right) mice with different diets. **k,** Quantification of differential signaling strength via differential analysis. **l,** Peak expression analysis of CM-EC communication, showing ligand-receptor pairs relative level (size) and intensity (color) according to genotypes and different diets. **m,** Immunofluorescence analysis of α-SMA^+^ or PDGFRα^+^ cells in CDH5*^mTmG^* mice with time-course of HDF + L-NAME treatment. n=3-6 per group. **n,** Immunofluorescence analysis of FBs (PDGFRα^+^ cells) and active ECs (αSMA^+^IB4^+^ cells) in Cre or cKO hearts after HFD + L-NAME treatment. n=3 per group. **o, p,** Immunofluorescence staining (o) and quantification (p) of human coronary artery endothelial cells (HCAECs) with conditioned medium from NRVMs. n=17 in Vehicle group, n=19 in TMP195 group, n=18 in PA + L-NAME group, n=29 in PA + L-NAME + TMP195 group. **q,** Working model. HDAC4-dependent mechanism in CMs affects ECs via ligand-receptor pairs of cardiocrine signaling. All the data are from three independent experiments. Data are mean ± s.e.m. **m, n,** two-way ANOVA with Sidak’s multiple-comparisons test. **p,** one-way ANOVA with Tukey’s multiple comparisons test. *P<0.05; **P<0.01; ****P<0.0001.

However, it seems unlikely that the profound phenotypic rescue, including the antifibrotic effects are not associated with gene expression changes in the affected cell types. Thus, we performed single nuclei (sn)RNA-seq to also analyze the non-cardiomyocyte cell populations. We chose the cKO model to avoid any effect of genetic manipulations on gene expression in other cell-types than cardiomyocytes. After rigorous quality control, a total of 203791 nuclei were retained for downstream analysis.

Unsupervised clustering via Uniform Manifold Approximation and Projection (UMAP) uncovered 12 distinct cellular profiles and did not show significant changes in distribution in different groups (Fig. 5e, f, Extended Data Fig. 7e). Pseudobulk DEG analysis revealed that the transcriptional impact of HFD + L-NAME treatment was most pronounced in endothelial cells (ECs), which exhibited the largest number of DEGs compared to fibroblasts (FBs) and cardiomyocytes (CMs). Notably, the vast majority of these transcriptomic alterations were found to be independent of *Hdac4* in cardiomyocytes although its deletion in cardiomyocytes prevents diastolic dysfuction (Fig. 5g, h). To further dissect the cellular-level heterogeneity and potential shifts after HFD + L-NAME treatment in hearts from Cre or cKO mice, we performed sub-clustering of the three major cell types (ECs, FBs and CMs). Unsupervised clustering and UMAP visualization identified 5 distinct EC subclusters (EC1-5), 8 FB subclusters (FB1-8), and 5 CM subclusters (CM1-5) (Fig. 5i). GO pathway enrichment analysis on these subtype-specific DEGs uncovered ECs were strongly and selectively enriched for terms related to GTPase activity, while FBs and CMs involved in the regulation of extracellular structure organization and glucokinase activity, respectively (Extended Data Fig. 7f-h).

To identify the specific cell populations driving the altered intercellular communication network in HFpEF, we analyzed the signaling shifts across major cell types and four distinct groups. Trajectory analysis of the aggregate cell-cell communication network revealed a universal upregulation of signaling interaction strengths in the HFpEF group. Notably, the endothelial lineage, comprising ECs and Lymphatic ECs (LECs), emerged as the most significantly altered populations, exhibiting the most substantial functional state transition across both sender and receiver roles this aberrant hyperconnectivity was significantly attenuated in cKO-HFpEF hearts (Fig. 5j). Quantification via differential analysis further confirmed that ECs and LECs displayed the highest net increase in both outgoing (sender) and incoming (receiver) signaling strengths (Fig. 5k). This result suggested that the profound remodeling of endothelial signaling in HFpEF is fundamentally driven by an HDAC4-dependent mechanism. Focusing specifically on cardiomyocyte-to-endothelial (CM-EC) communication, we identified ligand-receptor pairs that were significantly enriched in WT-HFpEF but suppressed in cKO-HFpEF hearts. Top-ranked *Hdac4*-dependent pairs included were Angpt2-Tek, Wnt5b-Fzd4, Gas6-Mertk and Vegfa-Vegfr1 (Fig. 5l), indicating that widespread endothelial activation acts as a central driver of the microenvironmental remodeling in HFpEF.

We validated this mechanism for ECs activation by lineage tracing experiments using CDH5*^mTmG^* mice that developed HFpEF. CDH5*^mTmG^* mice were treated with HFD + L-NAME for 3 days, 1 week, 5weeks and 7 weeks. HFpEF phenotypes were confirmed by measuring EF, E/e’ and MPI (Extended Data Fig. 8a-c) after treating with HFD + L-NAME for 7 weeks. By immunofluorescence analysis, we identified GFP stained ECs and cells that upon HFD + L-NAME overlap for α-SMA, providing confirmatory evidence for activation of ECs. However, we failed to identify an overlap with PDGFRα (Fig. 5m, Extended Data Fig. 9a, b), indicating that ECs do not transdifferentiate to FBs.

To investigate whether the activation of ECs is HDAC4-dependent, immunofluorescence experiments were performed by co-staining of α-SMA and the markers for ECs (IB4) or FBs (PDGFRα). We found evidence for substantial FB proliferation and EC activation in Cre but not cKO hearts (Fig. 5n). To mimic the two-hit HFpEF model *in vitro*, we treated neonatal rat ventricular myocytes (NRVMs) with palmitic acid (PA) and L-NAME. A robust increase of iNOS expression and LD accumulation was observed as signs for nitrosative and metabolic stress on a cellular level (Extended Data Fig. 10a, b). Taking advantage of this *in vitro* model, we found that human coronary artery endothelial cells (HCAECs) highly expressed α-SMA after exposure to the conditioned medium from NRVMs treated with PA+L-NAME, while this effect was reversed by conditioned medium from TMP195-pretreated NRVMs (Fig. 5o, p). Taking together, these data indicate that an HDAC4-dependent mechanism in CMs affects ECs possibly via ligand-receptor pairs of cardiocrine signaling (Fig. 5q).

## DISCUSSION

HFpEF is a multifactorial disease that lacks specific and efficient therapies, and in particular cardiomyocyte-directed therapies^31^. We studied preclinical animal models and patient samples to investigate the enzymatic activity class IIa HDACs in HFpEF^32^. HDAC4 as a member of the class IIa HDACs has much lower activity on standard acetyl-lysine substrates compared to a class IIa HDAC-specific-yet artificial-substrate (BOC-Lys(TFA)-AMC)^15–17,21^. Endogenous substrates of HDAC4 in muscle tissue have been proposed, but direct proof in conversion assays using recombinant proteins are missing and critical cellular pathways in cardiac diseases remain elusive^33^. Applying HDAC class-selective activity assays, we found that specifically the enzymatic activity of cardiac class IIa HDACs is increased in HFpEF.

A central finding of our study is the striking divergence between the effects of selective and non-selective HDAC inhibition in HFpEF. While pharmacological pan-HDAC inhibition resulted in mortality in mice with established cardiometabolic disease, selective inhibition of class IIa HDACs robustly reversed HFpEF phenotypes without overt adverse effects. This toxicity is consistent with prior reports demonstrating that class I HDACs particularly HDAC3 are essential for cardiac and systemic metabolic homeostasis, with genetic loss of HDAC3 causing lethality under high-fat diet conditions^34^. In line with this, we show that selective class I HDAC inhibition phenocopies the extensive increased chromatin accessibility observed with pan-HDAC inhibition, whereas selective class IIa HDAC inhibition did not. Together, these findings underscore that therapeutic efficacy and safety critically depend on HDAC class specificity, and that disruption of class I HDAC function likely underlies the detrimental effects of pan-HDAC inhibition in HFpEF.

Our findings integrate well with emerging therapeutic concepts in HFpEF. Dietary leucine supplementation has been reported to improve diastolic dysfunction in a rat model of HFpEF, which was associated with a reduced expression of *Hdac4*^35^. Recently, inhibition of a class IIb HDAC, HDAC6, has also been suggested to be effective to treat cardiometabolic HFpEF^36^ but it remains to be shown whether this relates specifically to HDAC6 since mice lacking *Hdac6* develop diastolic dysfunction^37^, and the underlying cell types have not yet been clearly identified.

We found that enzymatic activation of HDAC4 in cardiomyocytes causes diastolic dysfunction *in vivo* with EC dysfunction downstream of HDAC4. Emerging evidence suggested that crosstalk between CMs and ECs in a paracrine-dependent-pathway plays a crucial role in the development of several cardiovascular diseases^38^. However, the mechanism by which HDAC4 in CMs influence ECs function merits in depth investigations including the identification of the natural substrates of HDAC4 in cardiomyocytes. While established developmental models identify specific signaling axes, such as the NPPA-NPR3 interaction between atrial CMs and ECs, the factors driving this crosstalk in adult disease states are undefined^39^. We propose that enzymatic activation of HDAC4 does not act via gene expression control in CMs as it is well established for its non-enzymatic activity. In fact, histones seem not to be deacetylation targets of HDAC4, questioning the right nomenclature of this group of enzymes. Instead, we postulate that non-histone proteins need to be identified as enzymatic targets of HDAC4 in HFpEF. Given that nucleo-cytoplasmic shuttling is characteristic for HDAC4 upon oxidative stress and mitochondrial production of reactive oxygen species and dysfunction is closely associated with cardiometabolic HFpEF^26,40^, we propose to seek for targets in the cytoplasm.

In conclusion, our study demonstrates that enzymatic activation of HDAC4 drives HFpEF through non-epigenetic actions in cardiomyocytes, which leads through unexplored mechanisms to endothelial dysfunction and further microenvironmental remodeling. Pharmacological enzymatic inhibition by TMP195, a selective class IIa HDAC inhibitor, strongly reversed the observed cardiac HFpEF phenotypes. Thus, targeting the intrinsic enzymatic domain of HDAC4 is a promising translational approach to treat HFpEF. TMP195 as other selective class IIa HDAC inhibitors have not yet reached the clinical arena. Medicinal chemistry is ongoing to develop clinical candidates as first-in-class class IIa HDAC inhibitors to enable clinical trials validating the concept of this study in patients.

## Acknowledgments

We thank Jutta Krebs-Haupenthal, Silvia Harrack, Michaela Oestringer, Tanja Lüneburg and Joshua Hartmann for technical help. This work was supported by grants from Deutsche Forschungsgemeinschaft (DFG, German Research Foundation) (SFB1550, Project ID:464424253; Collaborative Research Center 1550, CRC1550 Molecular Circuits of Heart Disease; J. Backs, M. Dewenter, A. Schulze and R. Gilsbach); DZHK (Deutsches Zentrum fucr Herz-Kreislauf-Forschung-German Center for Cardiovascular Research) and BMBF (Bundesministerium fucr Bildung und Forschung) (J. Backs); Guangdong Basic and Applied Basic Research Foundation (Project ID: 2023A1515111097; J. Huang); China Scholarship Council (No. 202106380086; J. Fan). The authors acknowledge support by the state of Baden-Württemberg through bwHPC and the German Research Foundation (DFG) through grant INST 35/1597-1 FUGG.

## Author contributions

J.H. designed and conducted the experiments, performed the analyses, and wrote the first draft of the manuscript. F.S., J.F., Z.C., H.P., C.W., C.U.O., A.S., E.S., S.H., A.E., K.L., R.G., S.J., M.A.N. and M.D. generated tools, conducted additional experiments and analyzed data. F.S. generated *Hdac4^H968Y^* knock-in mice. N.G.C. provided advice and scientific interpretation of histological staining. R.A.B., I.F.P. and N.H. kindly provided samples from HFpEF mice, rats and patients biopsies, respectively. M.H. prepared the figures. M.D. and J.B. revised the manuscript. J.B. conceived the project and contributed to manuscript submission.

## Figure Legends

**Extended Data Fig. 1 | UMAP profiles and class IIa HDACs expression in different cell types. a-d,** *Hdac4* (a), *Hdac5* (b), *Hdac7* (c) and *Hdac9* (d) expression in different cell types from single-nucleus RNA sequencing analysis.

**Extended Data Fig. 2 | Cardiac and systemic phenotypes of Cre and cKO mice after 5-week with or without HFD + L-NAME treatment. a,** Weekly body weight of mice with different diets. n=6/ group. **b, c,** Tail-cuff systolic (b) and diastolic (c) blood pressure. n=6/ group. **d,** Blood glucose and area under the curve (AUC) of intraperitoneal glucose-tolerance test (iGTT). n=6/ group. **e,** Ratio of heart weight and tibia length. n=6/ group. **f,** Quantification of fluorescence intensity of adult cardiomyocytes (ACM) with bodipy staining. n=3 hearts/ group. Data are mean ± s.e.m. **a, d,** two-way ANOVA with Tukey’s multiple comparisons test. **b, c, e, j,** two-way ANOVA with Sidak’s multiple comparisons test. **f,** one-way ANOVA with Tukey’s multiple comparisons test. **P<0.01; ***P<0.001; ****P<0.0001.

**Extended Data Fig. 3** | **Cardiac and systemic phenotypes of *Hdac4^H968Y^* mice with HFD + L-NAME-induced HFpEF or TAC-induced HFrEF. a,** Strategy for generating *Hdac4^H968Y^* mouse line. sgRNAs were in red. Single-stranded oligodeoxynucleotides (ssODNs) were designed to provide not only the histidine to tyrosine (H968Y) mutation but also a RsaI restriction site (green). Additional mutations (*) were introduced by the template to mutate protospacer adjacent motif (PAM) sites (blue). **b,** The template ssODNs contained, next to the wanted mutation, an additional RsaI restriction site that served for genotyping purposes. **c,** Exemplary sequence trace of a heterozygous mouse. Mutated bases are indicated by a star (*). **d,** Weekly body weight of WT or H968Y mice with different diets. n=6/ group. **e, f,** Tail-cuff systolic (e) and diastolic (f) blood pressure. n=6/ group. **g,** Blood glucose and AUC of iGTT. n=6/ group. **h,** Ratio of heart weight and tibia length. n=6/ group. **i,** Representative images from echocardiographic pulsed wave Doppler (PWD), TD (tissue Doppler), P-V loop and bodipy staining in different groups. **j,** Representative EDPVR exponential curve during hemodynamic measurements in mice with or without HFD + L-NAME. **k,** Quantification of fluorescence intensity of ACM with Bodipy staining. n=3 hearts/ group. **l,** Stenotic velocity of transverse aortic artery of mice after TAC surgery. n=6/ group. **m-o,** Ratio of heart weight to tibia length (m), percentage of FS (n) and EF (o) of sham and TAC mice. FS, fractional shortening. n=4 in sham group. n=6 in TAC group. Data are mean ± s.e.m. **d, g,** two-way ANOVA with Tukey’s multiple comparisons test. **e, f, h, m-o,** two-way ANOVA with Sidak’s multiple comparisons test. **k,** one-way ANOVA with Tukey’s multiple comparisons test. **l,** two-tailed unpaired t-test. *P<0.05; **P<0.01; ***P<0.001; ****P<0.0001.

**Extended Data Fig. 4 | TMP195 selectively inhibits class IIa HDAC activity. a-c,** Class I, class IIa or class IIb HDAC activity of recombinant HDAC4 protein (a), induced pluripotent stem cells derived cardiomyocytes (iPSCs; b) and cardiac tissue (c) in response to different doses of TMP 195 treatment. **d-e,** Class I (d), class IIa (e) or class IIb (f) HDAC activity of various doses of proteins from cardiac tissue by *ex vivo* treating with or without TMP195. **g-i,** Class I (g), class IIa (h) or class IIb (i) HDAC activity of various doses of recHDAC4 by *in vitro* treated with or without TMP195. **j,** Cardiac class IIa HDAC activity after 4h injection of HDAC inhibitors. n=3/ group. Data are mean ± s.e.m. Statistical analysis was performed by one-way ANOVA with Tukey’s multiple comparisons test. *P<0.05.

**Extended Data Fig. 5 | Systemic and cardiac phenotypes of HFpEF mice with early intervention of HDAC inhibitors. a, b,** Systolic (a) and diastolic (b) blood pressure measured in a non-invasive way. n=6/ group except n=5 in HFD + L-NAME + DMSO group. **c,** Blood glucose and AUC of iGTT. n=6 in Ctrl Diet+ DMSO; n=5 in HFD + L-NAME + DMSO; n=5 in HFD + L-NAME + TMP195 group; n=6 in HFD + L-NAME + SAHA group. **d,** Representative images from pulsed wave Doppler (PWD), TD (tissue Doppler), and P-V loop in different groups. **e,** EDPVR exponential curve in indicated groups. **f,** *Col1a1/ Col3a1* mRNA expressions in cardiac samples from different groups. n=6 in Ctrl Diet + DMSO and HFD + L-NAME + SAHA group, n=5 in HFD + L-NAME + DMSO and HFD + L-NAME + TMP195 group. Data are mean ± s.e.m. **a-c, f,** one-way ANOVA with Tukey’s multiple comparisons test. *P<0.05; **P<0.01; ***P<0.001.

**Extended Data Fig. 6 | Systemic and cardiac phenotypes of different HFpEF mouse models with late intervention of HDAC inhibitors. a-c,** Non-invasive systolic (a), diastolic (b) blood pressure and the ratio between heart weight and tibia length (c) from indicated groups. n=4 in Ctrl Diet+ DMSO group; n=5 in HFD + L-NAME + DMSO; n=6 in HFD + L-NAME + TMP195 group. **d,** Representative images from PWD, TD and P-V loop in indicated groups. **e,** EDPVR exponential curve in different groups. **f,** Gene expressions from LV with different treatments. n=6 in Ctrl Diet + DMSO group; n=6∼7 in HFD + AngII + DMSO; n=7∼8 in HFD + AngII + TMP195 group. Data are mean ± s.e.m. Statistical analysis was performed by one-way ANOVA with Tukey’s multiple comparisons test. *P<0.05; **P<0.01; ***P<0.001; ****P<0.0001.

**Extended Data Fig. 7 | Differentially expressed genes analysis by RNA sequencing. a,** Volcano plot illustrating the significance of DEGs of CM nuclei in WT mice with or without HFD + L-NAME using p<0.05 and 1.5-fold regulation as cut-off criteria. **b,** Unsupervised clustering of DEGs of CM nuclei detected in Ctrl Diet versus HFD + L-NAME of WT mice, showing that genes and samples clustered according to HFD + L-NAME treatment. **c,** DEGs analysis of cardiac nuclei in WT or H968Y hearts after HFD + L-NAME treatment. **d,** Heatmap of ATAC-seq signal of HFD + L-NAME-responsive genes in WT and *H967Y* mutant. **e,** Distinct cellular profiles plotted by specific markers from single-nucleus RNA sequencing. **f-h,** GO pathway enrichment analysis on ECs (e), FBs (f) and CMs (g).

**Extended Data Fig. 8 | Cardiac phenotypes of CDH5*^mTmG^*mice with or without HFD + L-NAME treatment. a-c,** Percentage of EF (a), ratio between E wave and e’ wave (b) and MPI (c) from CDH5*^mTmG^* mice with different diets. n=6/ group. Data are mean ± s.e.m. Statistical analysis was done by two-tailed unpaired t-test. **P<0.01; ***P<0.001.

**Extended Data Fig. 9 | Immunofluorescence staining of CDH5*^mTmG^*mice with time-course of HFD + L-NAME treatment. a, b,** Representative images from immunofluorescence staining of α-SMA (a) and PDGFRα (b) in CDH5*^mTmG^* mice with time-course of HDF+L-NAME treatment.

**Extended Data Fig. 10 | Cellular stress model in NRVMs. a,** RNA expression of iNOS (Nos2) after 4h-treated with 0.1mM PA and 0.2mM L-NAME in NRVMs; n=3 per group. **b,** Representative images from Bodipy/DAPI staining of NRVMs with or without PA + L-NAME. Cells stained with DAPI were used as a negative control. Data are mean ± s.e.m. Statistical analysis was performed by two-tailed unpaired t-test. **P<0.01.

