## Supplemental Figures for "Selective class IIa HDAC inhibition reverses diastolic dysfunction in cardiometabolic HFpEF"

### Extended_Data_1

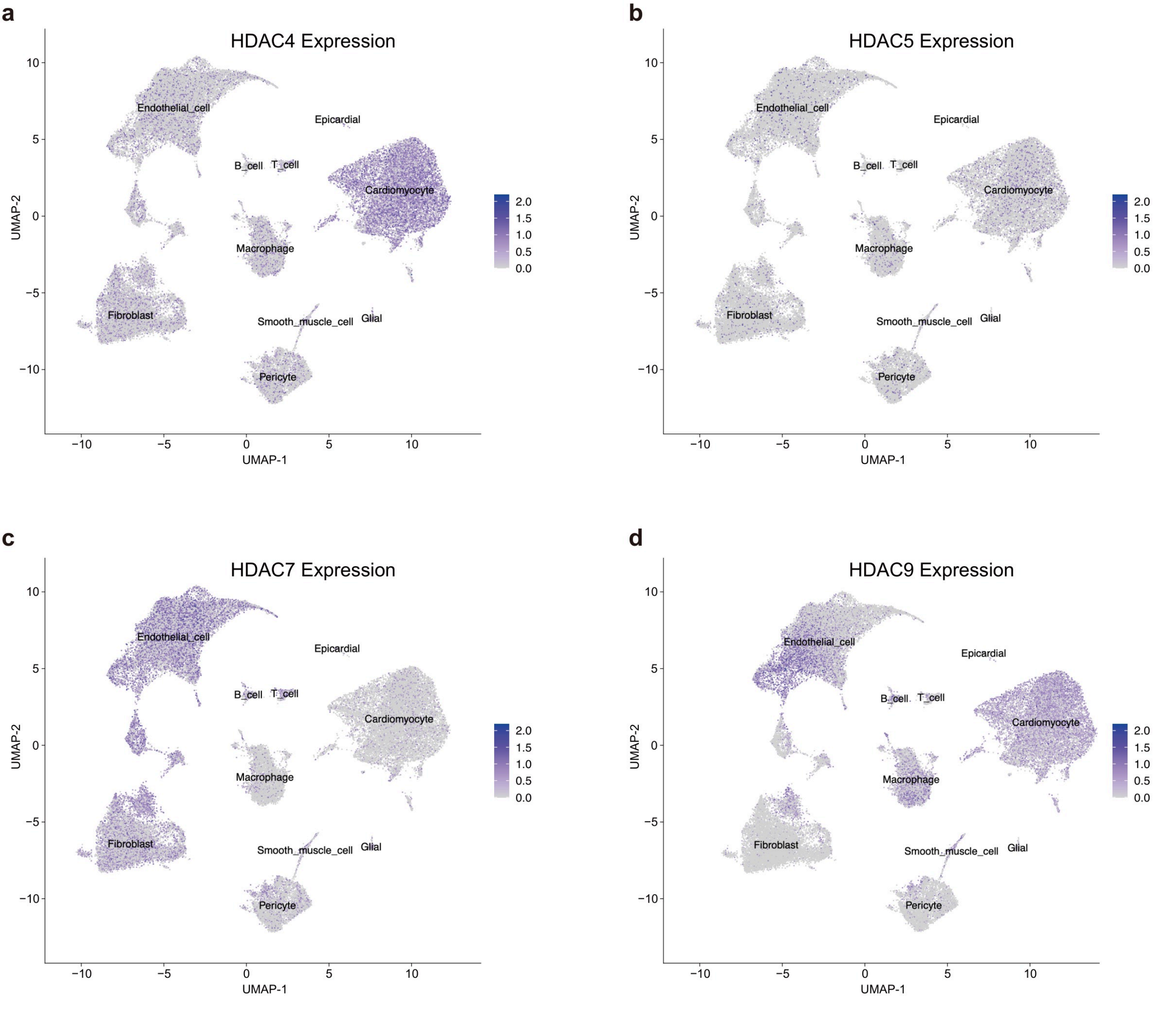

### Extended_Data_2

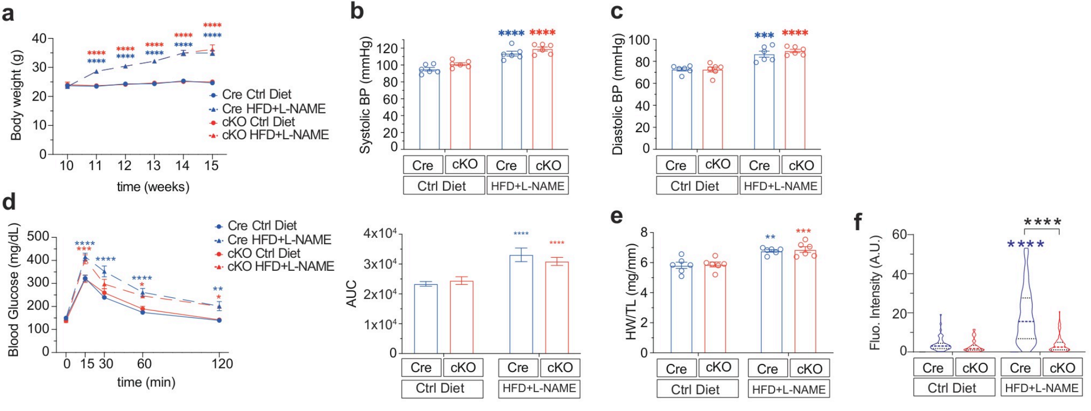

### Extended_Data_3

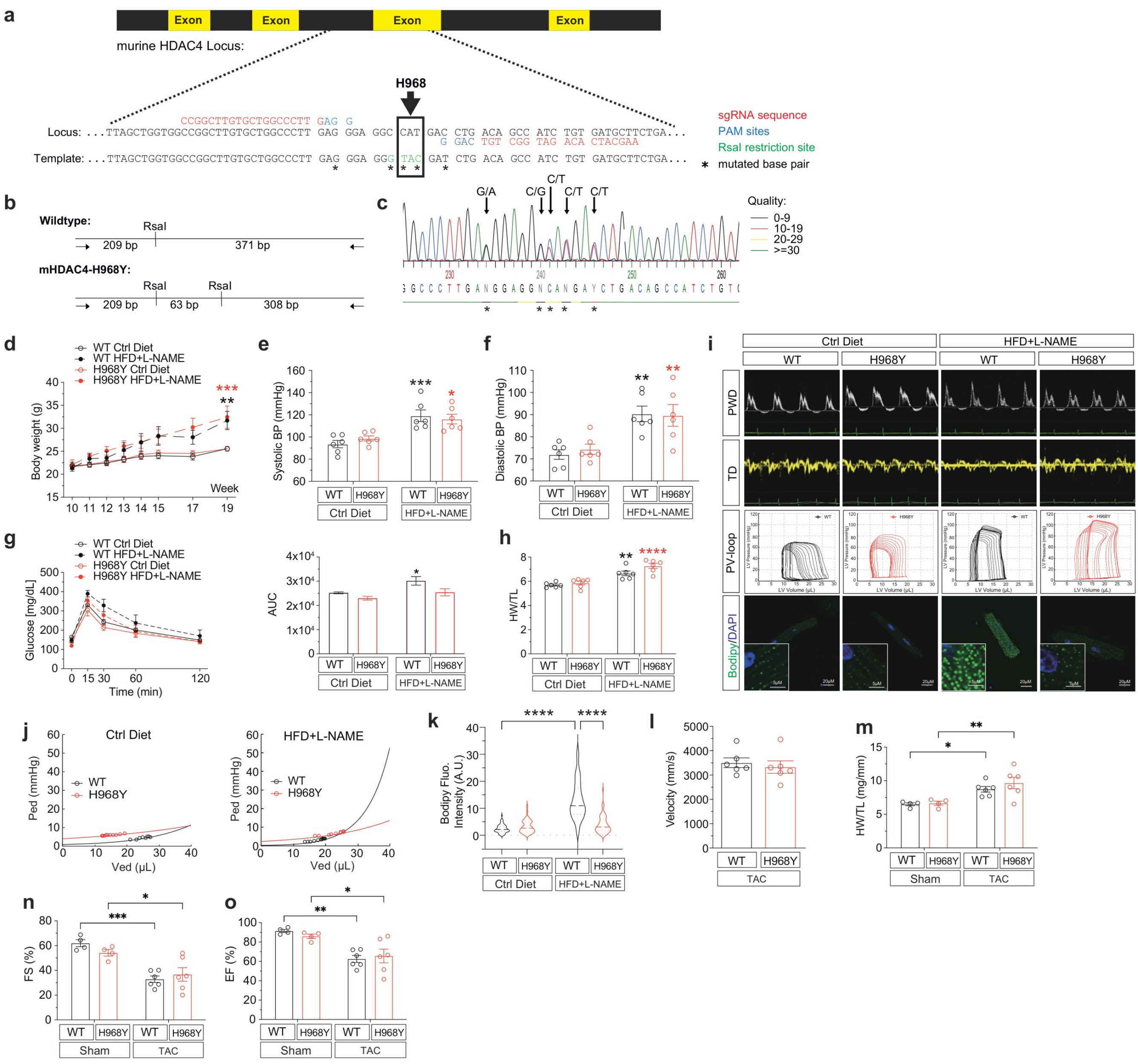

### Extended_Data_4

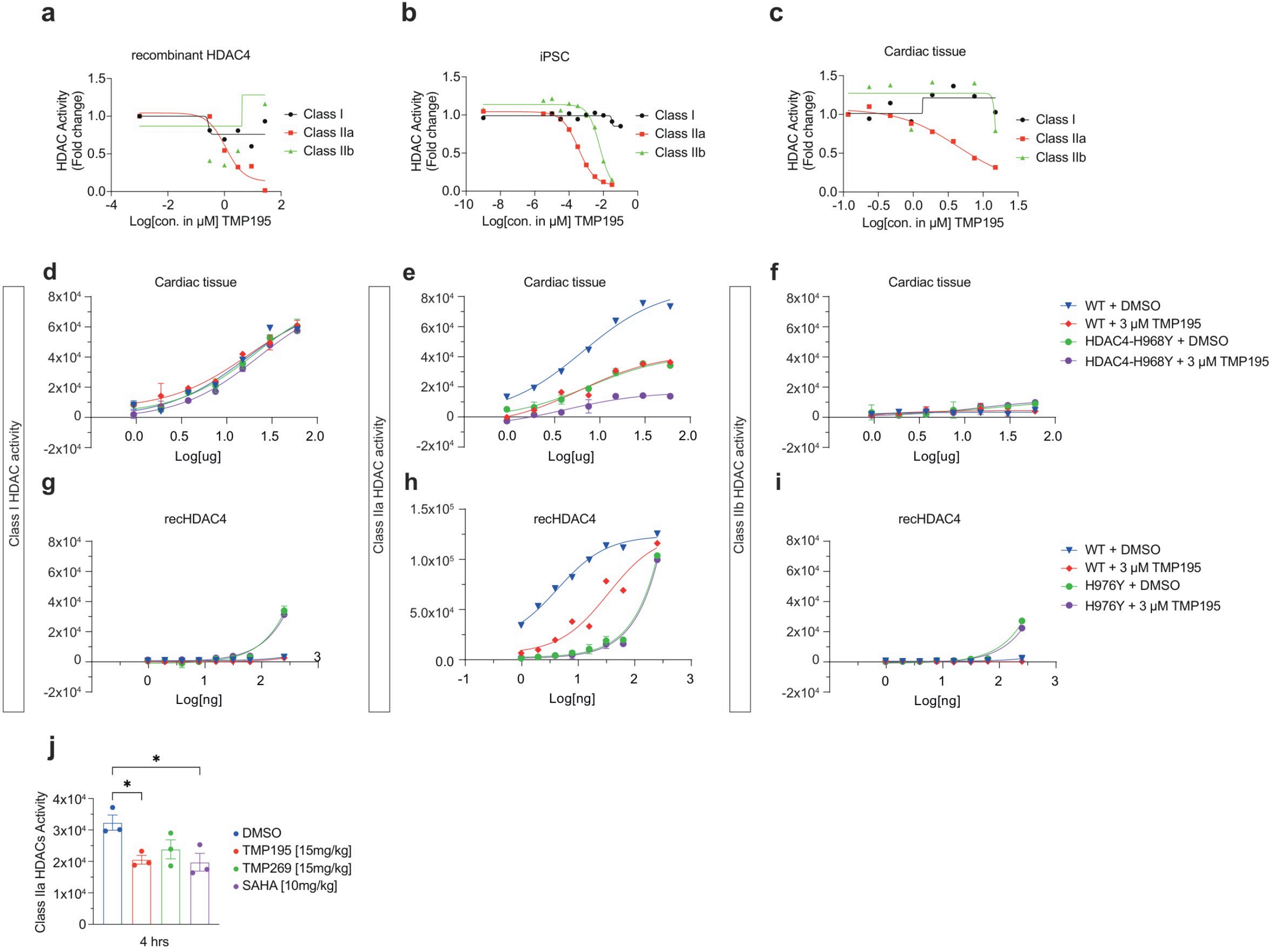

### Extended_Data_5

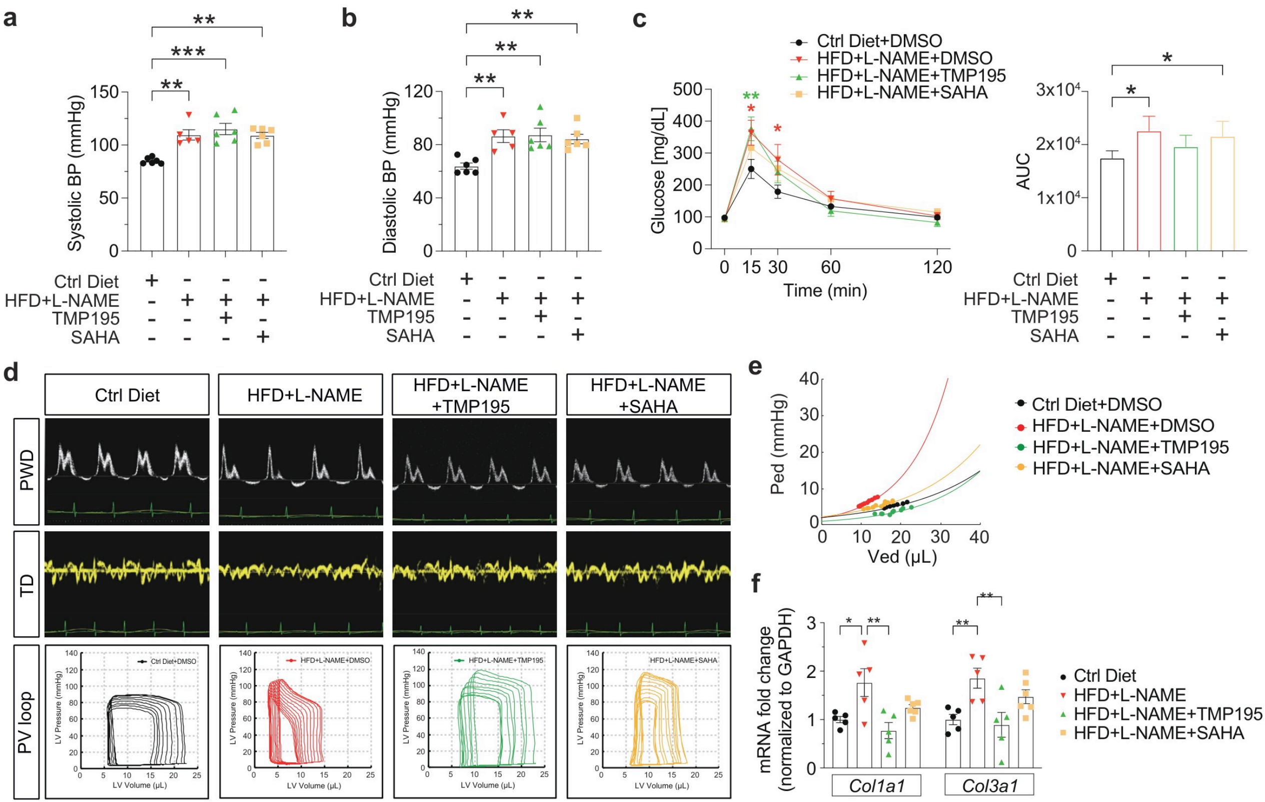

### Extended_Data_6

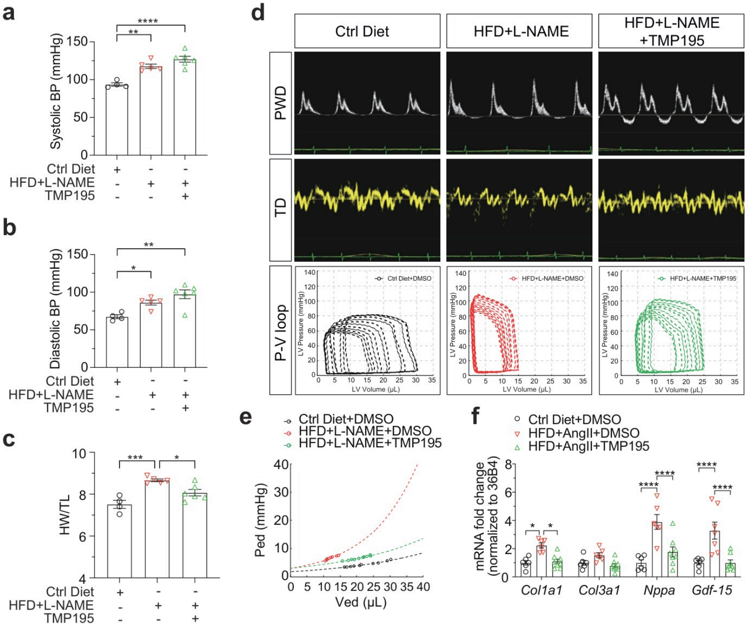

### Extended_Data_7

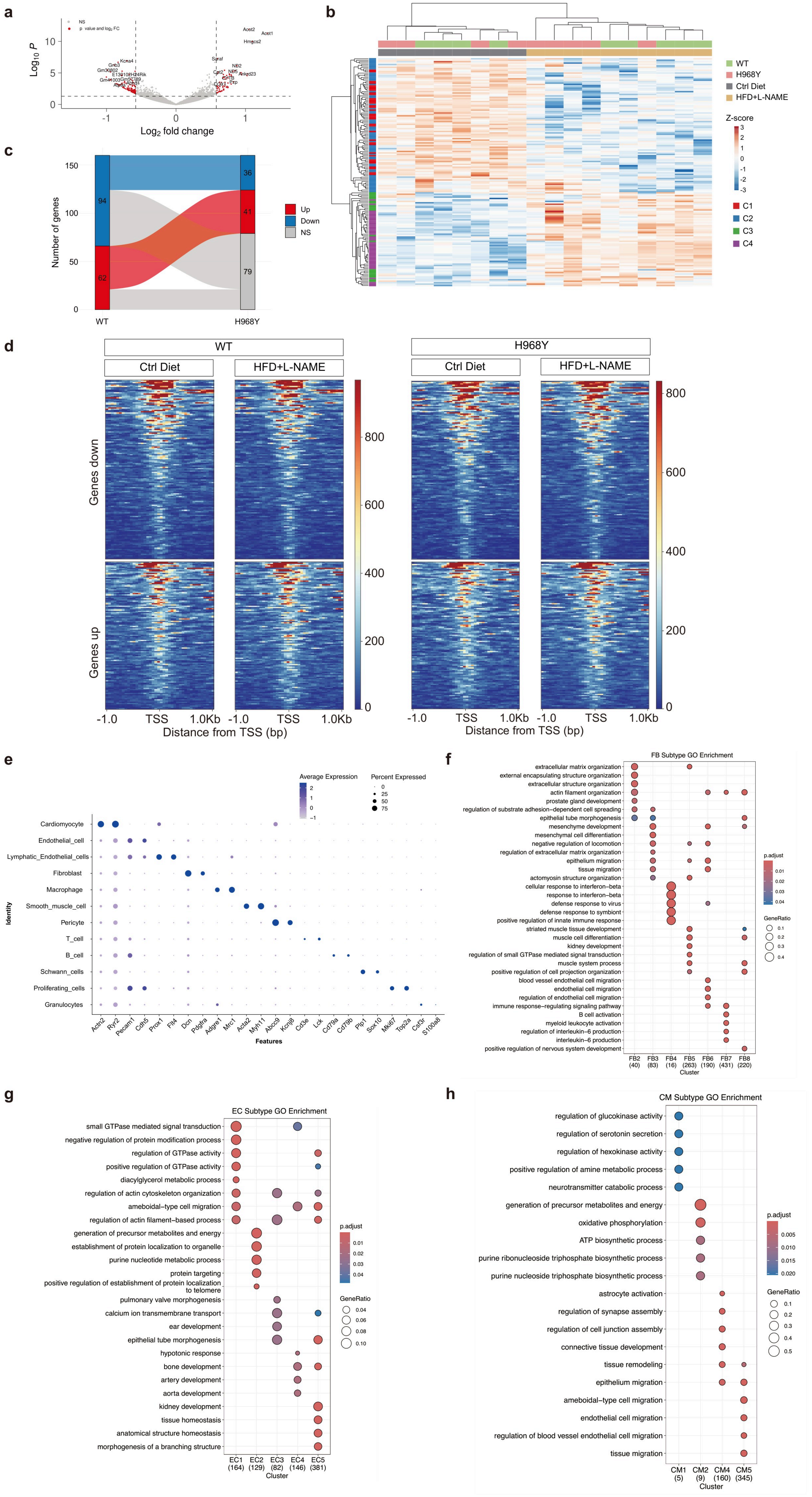

### Extended_Data_8

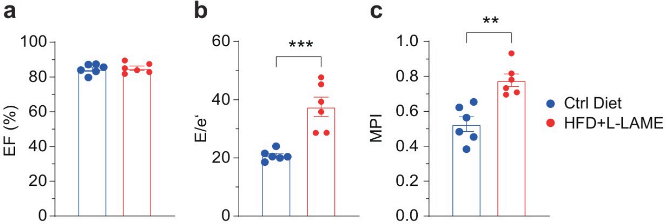

### Extended_Data_9

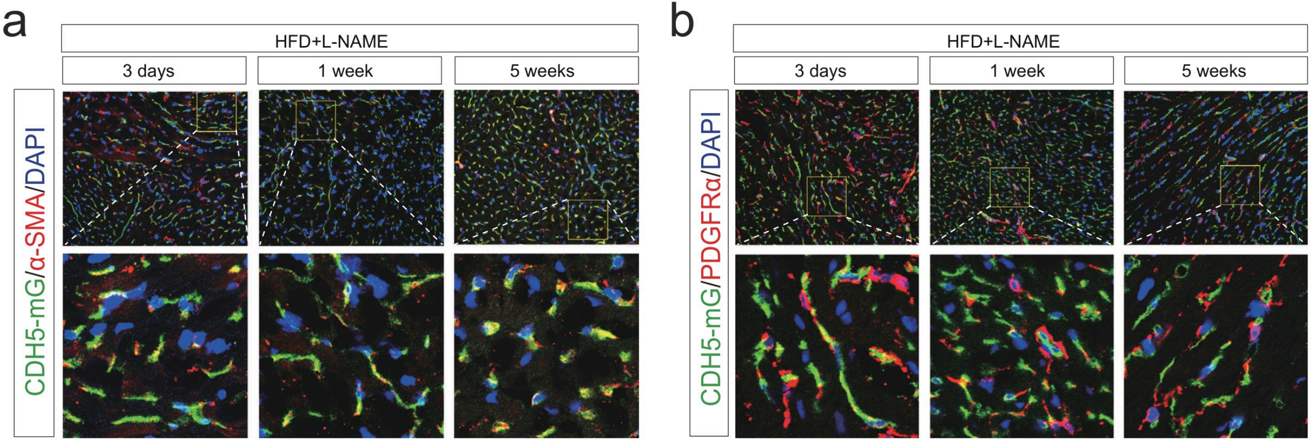

### Extended_Data_10

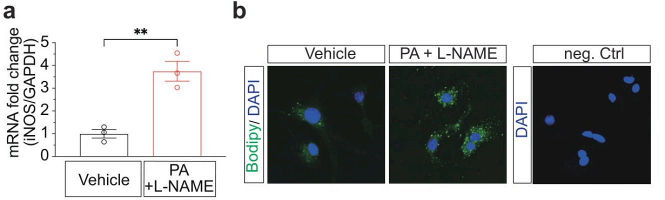
