## Supplementary material for "Selective class IIa HDAC inhibition reverses diastolic dysfunction in cardiometabolic HFpEF": Methods

#### Human myocardial samples

Myocardial biopsies were from patients with HFrEF or HFpEF, which were generously provided by Dr. Nazha Hamdani from Ruhr University Bochum, Germany.

#### Rat myocardial samples

Myocardial samples from ZSF1 obese (HFpEF) and ZSF1 lean (Ctrl) rats were provided by Dr. Ines Falcao-Pires in Faculty of Medicine of University of Porto, Portugal.

#### Animals

All experimental work of mice were approved by the Institutional Animal Care and Use Committee at the Regierungspräsidium Karlsruhe, Germany. As previous description[1], α-myosin heavy chain-Cre (α-MHC-Cre); *Hdac4*$^{loxp/loxp}$ mice with C57BL/6N background (*Hdac4*-cKO) and α-MHC-Cre positive littermates (Cre) were used for the experiments.

*Hdac4*$^{H968Y}$ mice were generated using CRISPR/Cas9. Following sgRNAs were chosen by using CRISPOR118: 5' AAGCATCACAGATGGCTGTC and 5' CCGGCTTGTGCTGGCCCTTG. T7 strings coding for sgRNAs were commercially provided by Thermo Scientific, which were used for in vitro transcription with Mega Short Script T7 Kit (Thermo Scientific) and Mega Clear Kit (Thermo Scientific) according to the manufacturer's protocol.

Injection Mix containing 3uM of a 120bp template ssODN (5' TTGACAAAACAGCTGATGGGCTTAGCTGGTGGCCGGCTTGTGCTGGCCCTT GAAGGAGGCTATGATCTGACAGCCATCTGTGATGCTTCTGAAGCCTGCGTG

TCTGCTCTGCTGGGAAAC, Eurofins genomics), 25ng/ul of each in vitro transcribed sgRNA, 25ng/ul Cas9 mRNA (Thermo Scientific) and 25ng/ul Cas9 protein (New Englad Biolabs) diluted in sterile-filtered injection buffer (5mM Tris, 0.1mM EDTA, pH 7.4) were injected into the cytoplasm of zygotes from super-ovulated female C57BL6/N mice. Zygotes were implanted into foster mice. Positive founder mice were bred back to C57BL6/N wildtype mice for six generations before they were used for experiments.

CDH5-CreERT2 mouse line was kindly provided by Prof. Ralf H. Adams. These mice were crossed to ROSA$^{mT/mG}$ reporter mice to mark ECs in a temporally controlled fashion as previously reported[2,3]. Cre activity was induced by tamoxifen solution (T5648, Sigma) via gavage.

Animal studies of aged female mice were approved by the Central Committee of Animal experiments (CCD) license number AVD105002016487 and the Animal Care and User Committee of the Groningen University (permit number 16487-07-07) and conducted in accordance with the ARRIVE guidelines[4], the general principles governing the use of animals in experiments of the European Communities (Directive 2010/63/EU) and Dutch legislation (The revised Experiments on Animals Act, 2014).

**HFD + L-NAME induced HFpEF mouse model**

Only male mice (10~12 weeks) were subjected to HFD + L-NAME exposure since previous report showed the potential protection in female mice[5]. Briefly, adult male mice were fed with HFD (60% kcal fat, E15742-347, Ssniff GmbH, Germany) or Ctrl Diet (13% kcal fat, E15748-047, Ssniff GmbH, Germany). A dose of 0.5g/L of L-NAME (N5751, Sigma) was supplied in drinking water with adjusted pH of 7.4.

**HFD+AngII induced HFpEF mouse model**

Aged female mice were fed a high-fat diet (HFD; Research Diets D17041409), or a low-fat equivalent chow (Crtl Diet; Research Diets D17041407) for 12 weeks. After 8 weeks of different diets, mice underwent surgery and an ALZET® osmotic mini pump (Model 2004) was implanted in a subcutaneous pocket on the back. For sham-treated mice, the subcutaneous pocket was closed without placement of a pump. During the last 4 weeks of the diet intervention, mice were infused with angiotensin II (AngII) (1.25 mg/kg/day) (Bachem) as previously described[6].

**Transverse aortic constriction**

Transverse aortic constriction (TAC) surgery was performed as previously descried[7]. Briefly, mice were anesthetized by maintaining 2% isoflurane after intubation. The transverse aortic artery was constricted with 7-0 suture around a 27-gauge blunt needle, which was immediately withdrawn after ligation. Mice in sham group were subjected to thoracotomy without ligation. All the mice with surgery were injected with Buprenorphine as a pain killer.

**HDAC inhibitors treatment**

TMP195 (HY-18361, Hölzel biotech), TMP269 (HY-18360, Hölzel biotech) and SAHA (SEL-S1047, Biozol) were dissolved in Vehicle (10% DMSO in corn oil). In HDACi dose finding experiment, doses of HDACi were indicated in different groups. For 4-week treatment of HDAC inhibitors, mice were treated with an intraperitoneally daily injection (i.p./q.d.) with total amount of 50 μL of TMP195 (15mg/kg/day), SAHA (10mg/kg/day) or Vehicle (10% DMSO).

**Animal Echocardiography and Doppler imaging**

Transthoracic echocardiographic measurements were performed using a Vevo 2100 system equipped with MX550D transducer (FUJIFILM VisualSonics), systolic function was evaluated in conscious by obtaining short views M-mode at midventricular level. Diastolic function was measured by pulsed wave and septal tissue doppler imaging at mitral valve level. Mice were maintained with 1~2% isoflurane during the measurement with heart rate above 400 beat per min as previous report[8]. Myocardial performance index (MPI) was calculated as the ratio of isovolumetric contraction time (IVCT) and isovolumetric relaxation time (IVRT) divided by ejection time (ET)[9]. All the data were measured at least 3 times and showed as mean values.

For aged female mice in HFD+AngII induced HFpEF model, 2D transthoracic echocardiographic measurements were performed using a Vevo 3100 system equipped with a 40-MHz MXX550D linear array transducer (FUJIFILM VisualSonics). Parasternal short and long axis views were obtained at Left Ventricular (LV) mid-papillary level under ~2% isoflurane maintaining. Vevo LAB software was used to assess cardiac dimensional and functional parameters and offline speckle tracking analysis (VEVO strain) was used to determine myocardial performance global longitudinal strain (GLS) as described[6].

**Tail-cuff blood pressure measurement**

Tail-cuff blood pressure was measured by CODA instrument (Kent Scientific). Briefly, mice were placed in the holders on a heating platform (37°C), blood pressure was recorded with a 15-cycle running set during stable conditions. Mean values were calculated from all accepted cycles.

**Glucose tolerance test**

Intraperitoneal glucose tolerance test (iGTT) was performed as described [10]. Briefly, mice were fasted for 6 hours before measuring baseline of tail blood glucose (as 0 min), then glucose (2 g/kg in saline) was intraperitoneally injected for glucose tolerance test. Blood glucose levels were recorded at 15, 30, 45 and 120 min after glucose injection. Aged mice were subjected to an oral glucose tolerance test (oGTT) in week 12. Mice were administered an oral bolus of glucose (2 g/kg) and blood glucose were measured in multiple time points.

**Exercise exhaustion test**

Mice were trained to adjusted to the uphill (20°) treadmill exercise at least three days. The fourth day, mice run with a speed of 16 cm/s for 5 min and then speed was increased by 3 cm/s every 2 min as previous descried[11]. The exhaustion of mice was defined as the inability to run on the treadmill for 10 sec despite mechanical prodding. Running distance was recorded once the mouse was exhausted.

**Hemodynamic pressure–volume measurement**

Mice were anesthetized by 5% isoflurane and then maintained with 1.5% isoflurane with ventilation, lack of response to firm pressure on hind paws was confirmed. Apical approach was used to obtain pressure-volume measurements by using mouse 1.4-F catheter (SPR 839, Millar Instruments) as described[12]. Load-independent parameters were obtained by inferior vena cava closure with a pause of ventilation. Data were analyzed by LabChart Pro v8.

**Palmitic acid solution**

Palmitic acid (PA; P0500, Sigma) was dissolved in a small volume of ethanol as previous protocol[13]. 10mM NaOH was added to convert it into palmitate solution, which was subsequently conjugated with 2% fatty acid free bovine serum albumin (BSA) in DMEM incomplete medium with a ratio of 2:1. After that, solution was in shaking water bath overnight at 55°C. PA solution was aliquot and stored at -20°C for future usage.

**HDAC activity assay**

HDAC activity assay was performed as previously described[14]. Briefly, tissue was lysed by lysis buffer with protease inhibitors (20mM Tris pH=7.4, 150mM NaCl, 1% NP-40, 1x Protease inhibitor cocktail). Protein concentrations of lysates were measured by BCA Protein Assay Kit (Thermo Scientific). 20~60μg of proteins from tissue lysates were used for HDAC activity measurement. For recombinant HDAC4 (rec HDAC4), 25ng of recombinant protein was used for activity assay. Substrates of class IIa (Boc-Lys(Tfa)-AMC, 4060676) and class IIb (Boc-Lys(Ac)-AMC, 4033972) were purchased form Bachem. Class I substrate (ZLPA, CBZ-Lys(Propionyl)-AMC) was a custom peptide from GeneScript. All the substrates were dissolved into DMSO with 50mM stock solutions. 5μL substrates with 1mM working solutions were added to 100μL protein lysates. After incubating at 37°C for 2.5 hours, 50μL/well of developer/stop solution (PBS with 1.5% Triton X-100, 3uM Trichostatin A, 0.75mg/mL Trypsin) was added and then incubated for 20min at 37°C. Particularly, to remove the signal of class I HDAC, 1μL of 300uM Apicidin (sc-202061, Santa Cruz) was previously incubated in

37°C for 30min before adding class IIb substrate. The 7-amino-4-methylcoumarin (AMC) fluorescence was measured by microplate reader (EnSpire™ Multimode Plate Reader, PerkinElmer, Inc.) with excitation/emission (Ex/Em) filters of 360/460 nm. Data were subtracted from background signals without protein.

For living cells HDAC activity measurement, cells were plated into 96-well with black bound. Culture mediums were removed and washed with pre-warm ADS buffer (116mM NaCl, 19.7mM HEPES, 9.4mM $NaH_2PO_4 \cdot 2H_2O$, 5.5mM Glucose, 5.4mM KCl, 0.8mM $MgSO_4 \cdot 7H_2O$) before treatment. Attached cells were incubated with ADS buffer for the whole treatment duration. NRVMs were treated with PA (0.1mM) and L-NAME (0.2mM) for 4 hours. N-Acetyl-L-cysteine (NAC, 1mM) was pre-incubated for 2 hours before challenging with PA+L-NAME. Class IIa HDAC substrate and developer/stop buffer were added at the last 2.5 hours and 20min of the experiments, respectively. The ACM fluorescence signal was measured as above. Background signals were from wells without cells.

**Expression plasmids and adenovirus**

*Hdac4*-WT and the mutant (H976Y) were cloned into pcDNA 3.1 Flag-tagged vector was generated by two PCR products (containing mutations and restriction enzyme site). Next, two PCR fragments are digested with appropriate enzymes and ligated to reassemble by in-frame coding sequence. All the generated plasmids were checked by restriction, PCR and sequencing.

For the generation of adenoviral infections, hHDAC4-WT was cloned into entry vector by BP reaction and then LR reaction was performed to transfer hHDAC4-WT into the pAd/CMV/V5-DEST. Further, these plasmids are digested by PAC-1 restriction enzyme, to expose ITRs. The Pac I-digested vector is transfected by lipofectamine 2000 in HEK293A cells to produce an adenoviral stock.

**Western Blot**

Lysates were diluted to an appropriate concentration and 6x Laemmli buffer (60 mM Tris, pH 6.8, 415 mM SDS, 50% (v/v) glycerol, 858 mM β-Mercaptoethanol, 0.1

mM Bromophenol Blue) was added. Proteins were separated on Tris-Glycine SDS-PAGE in running buffer and transferred to PVDF membrane in Pierce® Western Blot Transfer Buffer (Thermo Scientific, Waltham, Massachusetts). Depending on the requirements for each antibody, membranes were blocked in either 5% Skim Milk, 5% BSA or ROTI-Block (Carl ROTH, Karlsruhe, Germany) for at least 1 hour at room temperature, primary antibody was incubated in overnight at 4°C. Primary antibodies are as follows: Anti-Flag Antibody (MA1-91878, Thermo Fisher Scientific), GAPDH (MAB374, Merck Millipore), HDAC1 (#34589), HDAC2 (#57156) and HDAC3 (#85057) were purchased from Cell Signaling Technology, HDAC4 antibody was self-made.

Membranes were washed three times in TBST for 10 min and incubated in secondary antibody Goat anti-Mouse IgG-HRP (Cat. 1031-05, Southern Biotech) or Goat anti-rabbit IgG-HRP (Cat. 4050-05, Southern Biotech) at a 1:5000 dilution in TBST for 1 hour at room temperature. Eventually, membranes were incubated with chemiluminescent substrates (SuperSignal West Femto Maximum Sensitivity Substrate (Thermo Scientific, Waltham, Massachusetts, USA), Lumi-Light Western Blotting Substrate Plus (Merck Millipore, Burlington, Massachsetts, USA), Western Blot Luminol Reagent (Santa Cruz Biotechnology, Dallas, Texas, USA)) and imaged using a Vilber Fusion Fx System.

**Immunoprecipitation**

Cells were harvested from 6-well plates in IPA buffer (50 mM Tris, 150 mM NaCl, 1 mM EDTA, 1% Triton-X100) supplemented with SIGMAFAST Protease Inhibitor Cocktail Tablets (Sigma-Aldrich, St. Louis Missouri, USA) and Phosphatase Inhibitor Cocktail 2 (Sigma-Aldrich, St. Louis Missouri, USA). An equal amount from each sample was taken for input western blots and supplemented with 6x Laemmli buffer (60 mM Tris, pH 6.8, 415 mM SDS, 50% (v/v) glycerol, 858 mM β-Mercaptoethanol, 0.1 mM Bromophenol Blue). Equal amounts of the remaining samples were diluted to 500 µl with IPA buffer and incubated with Anti-Flag-beads (Sigma-Aldrich, St. Louis Missouri, USA) over night at 4°C on a rotator. Beads were prepared before as described in manufacturer's protocol. The next day beads were spun at 1,000 g for 1 min at 4°C. Supernatant was discarded and beads were washed three times in IPB buffer (50 mM Tris, 200 mM NaCl, 1 mM EDTA, 1% Triton-X100). Beads were washed once in 50 mM Tris. Supernatant was discarded and beads were resuspended in an appropriate volume 2x Laemmli without β-Mercaptoethanol. Samples were boiled for 5 min at 95°C. Supernatant was transferred to a new tube and supplemented with 6x Laemmli. Samples were boiled for another 5 min. Input and IP samples were used for Western Blot analysis.

**MEF2 luciferase assay**

Cells were transfected with plasmids encoding 3xMEF2-dependent reporter gene with or without Flag-*Hdac4*-WT or H976Y mutant for 24 hours. The luciferase assay was performed by Luciferase® Reporter Assay System kit (E1960, Promega) according to the manufacturer's instruction. The luciferase activity was detected by BMG Labtech FLUOstar OPTIMA luminometer.

**Expression and purification of recombinant protein**

The expression and purification of recombinant proteins were performed as previously described[7]. Recombinant HDAC4 WT and its mutant (H976Y) were performed using a baculoviral expression system. Briefly, human HDAC4 (HDAC4 WT) and its mutants were cloned into pFastBac HT A vector, which contained an N-terminal Tobacco Etch Virus (TEV)-cleavable hexahistidine (His6) tag. Recombinant bacmid was generated by electroporation of pFastBac HT His6–A-TEV- HDAC4 into *Escherichia coli* DH10EMBacY (Geneva Biotech, Genève, Switzerland). Recombinant bacmid was transfected into Sf9 cells (Thermo Fisher Scientific, Inc., Waltham, MA, USA) by X-tremeGENE™ HP DNA Transfection Reagent (Sigma-Aldrich, St. Louis, MO) to generate first generation virus (V0) culture. The V1 was used to infect Sf21 insect cells (~$0.9 \times 10^6$ cells/ ml) at a ratio of 1:100 in suspension culture using Sf-900 III serum-free medium (Life Technologies). Protein-expression procedures were performed at 27°C and 105 rpm for 72 h. The cells were collected by centrifuging at

600g and 4°C for 0.5 h and then resuspended in buffer A (50 mM Tris, pH=7.5, 250 mM NaCl, 20 mM imidazole, 10 mM, $MgCl_2$, 50 µM $ZnCl_2$, 1 mM DTT) with 7 U/ ml benzonase (70746, Novagen) and 1× cOmplete, EDTA-free protease inhibitor (11697498001, Roche). Insect-cell lysis was accomplished using a 25-ml Dounce homogenizer with 20 tractions on ice and cleared by ultracentrifugation (Optima XPN, Beckman Coulter, CA, USA) for 30 min at 45,000 rpm on Type 50.2 Ti rotor at 4°C.

Next, the cleared lysate was transferred on a 5-ml Protino Ni-NTA column (Machery-Nagel) which was pre-equilibrated with buffer A using Äkta Pure 25 system (GE Health Care). The bound protein was eluted with buffer B (50 mM Tris, pH=7.5, 250 mM NaCl, 250 mM imidazole, and 1 mM DTT) after extensive wash with buffer A. Elution fractions containing HDAC4 WT or its mutant were pooled, and the His6 tag was cleaved with TEV protease (produced in lab) during dialysis (50 mM Tris, pH 7.5, 250 mM NaCl and 250 mM imidazole, 1 mM DTT, 10 % glycerol) overnight at 4°C. In the following day, the protein was loaded again on a 5-ml Ni-NTA column to remove the TEV protease and the uncleaved proteins. The flow-through fractions containing the HDAC4 WT or its mutant were collected, pooled together, and brought to a volume of approximately 2 ml and further subjected to size exclusion chromatography using HiLoad 16/ 600 Superdex 200 PG column (GE healthcare) with 50 mM Tris pH=7.5, 250 mM NaCl, 1 mM DTT and 10% glycerol. The protein fractions were finally concentrated, snap-frozen in liquid nitrogen and stored at -80°C. The purity of the protein was evaluated by SDS–PAGE and subsequent Coomassie staining.

**Cardiomyocyte isolation**

Adult mouse cardiomyocytes (ACMs) were isolated from littermates of different groups by using Langendorff perfusion system as previous protocol[15].Briefly, hearts were removed and immediately perfused with pre-warm perfusion buffer (113mM NaCl, 4.7mM KCl, 0.6mM $KH_2PO_4$, 0.6mM $Na_2HPO_4$•$2H_2O$, 1.2mM $MgSO_4$•$7H_2O$, 12mM $NaHCO_3$, 10mM $KHCO_3$, 10mM HEPES, 30mM Taurine, 5.5mM glucose and 10mM BDM). After 1 min perfusion, individual heart was digested by 30mL digestion buffer (perfusion buffer containing 12.5μM $Ca^{2+}$, 0.042mg/mL liberase and 0.025% trypsin) for 10 min. The left ventricle was cut into small pieces and immediately added 2.5mL buffer with 1% BSA and 50μM $Ca^{2+}$. After 3 min gentle mix, mixture was filtered. ACMs were sedimentary by gravity. 5mL buffer with 0.5% BSA and 38μM $Ca^{2+}$ was added into cell pellet after removing the supernatant. Calcium treatment was performed for 5 steps from 62μM to 960μM in every 4 min. ACMs were plated in laminin-coated glass coverslips for staining.

Neonatal Rat Ventricular Myocytes (NRVMs) were isolated from newborn rats (1 to 3 days) according to the manufacturer instructions from Neonatal Heart Dissociation Kit (130-098-373, Miltenyi Biotec) and Neonatal Cardiomyocytes isolation kit (130-105-420, Miltenyi Biotec). NRVMs were cultured in DMEM with 10% FCS and 1% penicillin–streptomycin.

**Histological analysis and staining**

LV was cut transversally for two-chamber view and fixed in 4 % paraformaldehyde (PFA) in PBS before paraffin embedding. Samples were cut into 4μm thick sections and stained with Masson's trichrome as previously performed[16]. Images were obtained by Olympus BX51 microscope (Olympus America, Inc.).

For cryo-sections, samples were dehydrated in a sucrose gradient and embedded in O.C.T. solution (Tissue-Tek) after fixation with 4 % PFA as previously described[17]. α-SMA (ab32575, Abcam), PDGFRα (AF1062, R&D), Isolectin GS-IB4 (I21414, Invitrogen) were used as primary antibodies. Alexa Fluor 488 donkey anti-rabbit (A21206), Alexa Fluor 594 goat anti-rabbit (A11012), Alexa Fluor 647 donkey anti-goat (A21447) and DAPI were purchased from Life Technologies. Texas Red Streptavidin (SA-5006) was purchased from Vector Laboratories.

NRVMs or ACMs were plated on laminin-coated glass coverslips and fixed with 4% PFA. BODIPY™ 493/503 (D3922, Invitrogen) was used at a dilution of 1:1000 in 0.1% Triton X-100 for 1 hour at room temperature. Nuclei was stained by DAPI at a dilution of 1:5000.

Slides were mounted by fluorescence mounting medium (S3023, Agilent) and visualized by a confocal microscope (Leica SP8) and AxioObserver (Zeiss). Image quantification was performed by ImageJ software.

**Quantitative real time polymerase chain reaction (qPCR)**

Total RNA was extracted with TRIzol reagent and 500~1000ng of RNA was reverse transcribed to cDNA using the QuantiTect RT kit (Qiagen) or cDNA synthesis kit (Thermo Scientific). Gene expression levels were performed by qPCR analyses by using a SYBR green super mix. Relative quantification was performed to evaluate target genes expression by normalizing to GAPDH or 36B4. Primers sequences were displayed as follows (forward, reverse sequences, respectively): Mouse *Gapdh*: FCGTGCCGCCTGGAGAAACC, TGGAAGAGTGGGAGTTGCTGTTG; Mouse *36b4*: AAGCGCGTCCTGGCATTGTC, GCAGCCGCAAATGCAGATGG; Mouse *Col1a1*: AGAGCATGACCGATGGATTC, CGCTGTTCTTGCAGTGATAG; Mouse *Col3a1*: ACGTAAGCACTGGTGGACAG, CAGGAGGGCCATAGCTGAAC; Mouse *Nppa*: GCTTCCAGGCCATATTGGAG, GGTGGTCTAGCAGGTTCTTG; Mouse *Gdf-15*: TGACCCAGCTGTCCGGATAC, GTGCACGCGGTAGGCTTC. Rat *Gapdh*: GGTGGACCTCATGGCCTACA, CTCTCTTGCTCTCAGTATCCTTGCT; Rat *Nos2*: GCATCGGCAGGATTCAGTGG, GGAACACAGTAATGGCCG.

**Human induced pluripotent stem cells and cardiomyocyte differentiation**

For human induced pluripotent stem cells (hiPSC)-cardiomyocyte differentiation, hiPSC (https://hpscreg.eu/cell-line/RUCDRi002-A) were cultured in StemMACS Brew (Milteny Biotec) on Matrigel (Growth Factor Reduced, Corning) coated plates, mesoderm was induced by 4 μmol/L CHIR (TargetMol) for 24 h, followed by 2 μmol CHIR for additional 24 h in base medium (RPMI1640 (Gibco) supplemented with 213 μg/mL L-ascorbic acid (abcr GmbH) and 500 μg/mL human serum albumin (PanBiotech)). Cardiac mesoderm was induced by 5 μmol/L IWP2 (Selleckchem) in base medium for 2 days and cells were differentiated to cardiomyocytes in base medium for 4 days. Cardiomyocytes were cultured in RPMI1640 supplemented with B27

(Themo Fisher Scientific) before metabolic selection with RPMI no glucose (Gibco) with 2.2 mmol/L Lactate (Sigma Aldrich). Cells were detached with 10x TrypLE (Gibco) and 50,000 hiPSC-cardiomyocytes were seeded in a 96-multiwell plate and cultured for 2 weeks in RPMI1640 supplemented with B27 before treatment with Trichostatin A (TSA) (10 µmol/L final concentration, Sigma Aldrich), Romidepsin (300 nmol/L final concentration, Hycultec), TMP195 (300 nmol/L final concentration, MedChemExpress). DMSO treated hiPSC-cardiomyocytes served as control. Cells were harvested 3 h and further processed for ATAC sequencing.

**Cardiomyocyte nuclei isolation**

CM nuclei were isolated from the left ventricles of WT and H968Y mice following 5 weeks of HFD + L-NAME or control diets using fluorescence-activated nuclei sorting (FANS) as previously reported[18-20]. Draq-7 (Invitrogen, D15106) was used to stain all cardiac nuclei. CM nuclei were stained with rabbit anti-PCM-1 antibody (1:1000, Sigma-Aldrich, HPA023374) and mouse anti-PLN antibody (1:1000, Badrilla, A010-14). Anti-rabbit antibody conjugated to Alexa 568 (1:1000, Life Technologies, A11011) and anti-mouse antibody conjugated to Alexa 488 (1:1000, Life Technologies, A11029) were used as secondary antibodies. CM nuclei ($PCM1^+$, $PLN^+$ and Draq-$7^+$) were sorted using a SH800S cell sorter (Sony Biotechnology).

**RNA-seq library preparation of CM nuclei**

RNA was isolated using the RNeasy Micro Kit (Qiagen, 74004), with on-column DNase treatment. RNA-seq libraries were generated using SMART-Seq Total RNA Pico Input RNA-seq kit (ZapR Mammalian with UMIs, Takara, 634356). Libraries were sequenced in paired-end mode using an AVITI sequencer (Element Biosciences) using AVITI 2x75 Sequencing Kit Cloudbreak FS (860-00015).

RNA STAR (v2.7.10)[21] was used for mapping to mm39 genome. UMIs were extracted and used for deduplication using UMI-tools (v1.1.6)[22]. Gene counts were determined using featureCounts (v2.1.1)[23]. Differential gene expression was calculated using DESeq2 (v2.11.40.8)[24]. Statistical significance threshold was assigned at P value

<0.05 and abs(fold change)>1.5.

**Assay for transposase-accessible chromatin sequencing (ATAC-seq)**

CM nuclei were isolated from the left ventricles of WT and H968Y mice with HFD + L-NAME or Ctrl Diet feeding using the FANS method[18-20] and OMNI-ATAC-seq was performed[25]. Libraries were sequenced in paired-end mode using AVITI sequencer (Element Biosciences) using AVITI 2x75 Sequencing Kit Cloudbreak FS (860-00015). Adapters were trimmed using Cutadapt (v1.16.5). Reads were mapped to mm39 genome using Bowtie2 (v2.4.2)[26] in paired-end mode. Duplicated reads were identified using MarkDuplicates (v2.18.2.2) (http://broadinstitute.github.io/picard/). Coverage plots were generated using bamCoverage (v3.5.4)[27].

For hiPSC ATAC sequencing, cells were lysed in Lysis Buffer (10 mM TRIS-HCl, pH7.5, 10 mM NaCl, 3 mM MgCl2, 0.1% v/v NP-40, 0.1% v/v Tween-20, 0.01% v/v Digitonin) for 3 min on ice and nuclei were isolated by centrifugation at 1000 xg for 5 min, 4 °C. Tagmentation mix (25 µL 2x Tagment Buffer, 0.5 µL 10% Tween-20, 0.25 µL 2% Digitonin 2.5 µL Tn5 Transposase) was added to nuclei (50 µL per total reaction mix) and incubated for 30 min at 37 °C shaking. DNA was isolated (QiAquick, Qiagen) and library was prepared using NEBNext High-Fidelity 2x PCR MasterMix (NEB). Library was purified using AMPure XP Beads (Beckman Coulter). Library quality was assessed by Qubit fluorometric quantification (Thermo Fisher Scientific) and qPCR as well as by Agilent High Sensitivity DNA Bioanalysis on a TapeStation (Agilent). Adapters were added using theAdept Rapid PCR-Plus Protocol according to manufacturer's protocol (Element Biosciences). Sequencing was performed on an Element AVITI System Sequencing Instrument (Element Biosciences). Paired-end reads were aligned to the human reference genome (hg38) using Bowtie2, and aligned reads were stored as BAM files. Peaks were called using MACS2 with default parameters. Differential chromatin accessibility was assessed using the DiffBind R package. Initially, all BAM and corresponding BED peak files were loaded into a DiffBind object, and read counts for all peaks were computed across all samples using the dba.count function. Pairwise contrasts were defined between all conditions, and differential analysis for the TSA versus DMSO group contrast was performed with DESeq2 through dba.analyze. All differentially accessible peaks were selected based on the lowest false discovery rate (FDR) from the resulting differential report (dba.report), which provided fold-change and statistical significance values . To include all conditions in the visualization, the corresponding counts for these peaks were retrieved from the full count matrix using dba.peakset (bRetrieve = TRUE). Counts were log2-transformed (log2(counts + 1)) and standardized on a per-peak basis (row-wise Z-score) to allow comparison across samples. Heatmaps were generated with the pheatmap R package, clustering both rows (peaks) and columns (samples). This approach allowed visualization of the top differentially accessible loci between DMSO and TSA treated hiPSC-cardiomyocytes, while simultaneously comparing their accessibility patterns across all other experimental groups.

**Nuclei isolation and single nuclear RNA sequencing analysis**

Left ventricular tissues from cKO and Cre control mice (1-week HFD + L-NAME or Ctrl Diet) were dissociated using a gentleMACS (Miltenyi Biotec) as previously reported[28]. Briefly, homogenates were 30-μm filtered, centrifuged, and nuclei liberated in lysis buffer. Fluorescence-activated nuclei sorting (FANS) purified nuclei were processed via Chromium iX (10× Genomics) and sequenced on the Element AVITI System. Reads were aligned to mm10 using Cell Ranger (v7.1.0)[29]. Downstream analysis was performed in R using Seurat (v5.1.0)[30]. For quality control (QC), raw gene expression matrices were imported, and Nuclei meeting the following criteria were retained: > 300 detected genes per cell; > 1,000 unique molecular identifiers (UMIs); a $\log_{10}$ GenesPerUMI score > 0.8 to ensure library complexity and < 0.4 mitochondrial content. Following QC, ambient RNA was removed using celda (v1.18.2), which was from Campbell J, Yang S, Wang Z, Corbett S, Koga Y (2025). celda: CEllular Latent Dirichlet Allocation. doi:10.18129/B9.bioc.celda, R package version 1.26.0, https://bioconductor.org/packages/celda. R package, and potential doublets were identified and removed using DoubletFinder[31] (v2.0.4) R package. All furthering data integration and normalization were followed by the Seurat tutorial protocol. Cell clusters were annotated based on the expression of canonical marker genes. The same integration and clustering pipeline were applied to identify specific subtypes.

**Differential expression and functional enrichment analysis**

To identify differentially expressed genes (DEGs), we employed a pseudobulk strategy by aggregating single-cell counts to the biological replicate level using the *AggregateExpression* function in Seurat. Statistical differential analysis was performed using DESeq2[24] (v1.42.0) R package, with significance defined as a nominal p-value < 0.01, $|\log_2\text{Fold Change}| > 0.25$. Gene Othology (GO) enrichment analysis was performed using clusterProfiler[32] (v4.10.0) R package.

**Cell-cell communication analysis and visualization**

Cell-cell communication networks were inferred using CellChat[33] (v2.1.2) R package. The analysis was restricted to major cell types selected based on the relatively abundance: cardiomyocytes, endothelial cells, lymphatic endothelial cells, fibroblasts, macrophages, pericytes, and smooth muscle cells. The ligand-receptor interactions were modeled using the CellChatDB mouse database, specifically filtering for "Secreted Signaling" and "Cell-Cell Contact" categories to focus on biologically relevant interactions based on the rationale. The inferred networks and interaction strengths were generated using CellChat and visualized by ggplot2 (v3.5.1), which was by Wickham H (2016). ggplot2: Elegant Graphics for Data Analysis. Springer-Verlag New York. ISBN 978-3-319-24277-4, https://ggplot2.tidyverse.org. R package.

**Statistical analysis**

All the data were presented as mean ± standard errors of the mean (SEM). Significance of experiments with two groups was analyzed by two-tailed unpaired Student's t-test. The $\chi^2$-statistic was used for comparison of proportions as appropriate. One-way plus Tukey's post hoc test or two-way ANOVA with Sidak's post hoc test was used for multiple comparisons where appropriate in experiments with more than two groups. All statistical analyses were performed by GraphPad Prism Software (version 9), SPSS (version 23) and R Studio (version 4.2.3). A p-value of <0.05 was considered statistical significance.

### Data availability

The authors declare that the data supporting the findings of this study are available within the paper and its supplementary information. Source Data of the manuscript can be obtained from the corresponding author upon request.
